# Design and validation of the first family of photo-activatable ligands for melatonin receptors

**DOI:** 10.1101/2022.05.05.490738

**Authors:** Gloria Somalo-Barranco, Carme Serra, David Lyons, Hugh D. Piggins, Ralf Jockers, Amadeu Llebaria

## Abstract

Melatonin is a neurohormone released in a circadian manner with peak levels during the night. In mammals, melatonin mediates its effects mainly through G protein-coupled MT_1_ and MT_2_ receptors. Drugs acting on melatonin receptors are indicated for circadian rhythm- and sleep-related disorders and major depression. Pharmacological tools to study the activation of these receptors with high temporal resolution are lacking. Here, we synthesized a family of light-activatable caged melatonin compounds by covalently attaching *o*-nitrobenzyl (*o-*NB) or coumarin photocleavable groups to the *N-*position of melatonin. All caged compounds showed the expected decrease in binding affinity for MT_1_ and MT_2_. Among them, the *o-*NB derivative MCS-0382 showed the best uncaging and biological properties upon light activation with a significant increase (2.5 log left-shift) in affinity and potency in melatonin receptor binding and signaling, respectively. Generation of biologically active melatonin from MCS-0382 was further demonstrated by its ability to modulate the excitation of SCN neurons in rat brain slices. MCS-0382 is now available to study melatonin effects in a temporally controlled manner in cellular and physiological settings.

## INTRODUCTION

Melatonin (5-methoxy-*N*-acetyltryptamine) is a hormone that is predominantly secreted by the pineal gland, following a circadian rhythm with high levels at night.^1^ Melatonin regulates many physiological functions in the body including the regulation of biological rhythms, sleep, pain, retinal, neuronal, and immune functions.^2^ These effects are mainly mediated through the activation of two members of the G protein-coupled receptor (GPCR) family, MT_1_ and MT_2_ receptors, which preferentially couple to Gi/o proteins.^3^ Commercialized drugs targeting melatonin receptors are used to ameliorate sleep onset, depressive disorders and dysfunctions related to the circadian rhythms.^2^ Future therapeutic applications include the treatment of pain, inflammation, immune system disorders, and metabolic and neurodegenerative diseases.^2^

Light is now increasingly recognized as an ideal external control element to modulate physiological systems with a high spatiotemporal precision associated to a low toxicity and high safety.^4^ In this context, photopharmacology is presented as a discipline based on the development of molecular probes whose biological activity can be regulated by light. Light can be adjusted to most cellular processes, not interfering or damaging the living system and it can be manipulated remotely. Furthermore, the extent of its effect can be precisely regulated, through the adjustment of the wavelength and intensity of the light.^5^ Two of the most widely used molecular approaches that incorporate light to control biological processes are photoswitches and caged compounds, whose main difference is the reversibility of the process upon the application of light.^6^ Caged compounds are light-sensitive probes that are obtained by attaching covalently a photo-cleavable group to a bioactive molecule. This modification is designed to render the new ligand inactive to the target receptor. Upon suitable light conditions, an irreversible photolytic reaction is triggered in the caged molecule, which releases specifically the active molecule at the action site.^7^

Light-dependent control of receptor activity has been already achieved for several GPCRs,^8,9^ contributing to a better understanding of receptor function. In particular, light-controlled release of the active ligand upon uncaging provide valuable insights about receptor-induced signaling or binding kinetics.^10^ Photopharmacology has never been applied to melatonin receptors but would be very useful tools for precise spatial-temporal activation of melatonin receptor in vitro and in vivo. Using the well-known photo-cleavable groups *o*-nitrobenzyl (*o-*NB) and coumarin derivatives, we have developed here the first family of caged melatonin ligands, with the objective to modulate the activation of melatonin receptors in a light-controlled manner in vitro and in vivo.

## RESULTS AND DISCUSSION

### Design and chemical synthesis of caged melatonin compounds

A successful design of a caged compound is based in the attachment of a light-sensitive moiety or photocleavable-protecting group (PPG) in a region of the active molecule that is relevant for ligand recognition and activation of the target receptor.^7,11^ Our initial objective was to design a caged derivative of melatonin suitable to release this biomolecule upon illumination in biological environments. Structure-activity relationship (SAR) studies on melatonin receptors revealed the importance of the methoxy substituent at the C5 position of melatonin and the alkylamide side chain on the intrinsic activity and high binding affinity to its receptors.^1,12^ However, selective attachment of the PPG to the acetamide side chain was not considered initially a good approach, due to the recognized difficulties in amide photorelease.^13^ In spite that some solutions have been found for caging primary amides,^14–17^ no reports support its use in the release of secondary amides such as melatonin. Moreover, attachment of the PPG at the C5 methoxy was also discarded, due to the inexistence of chemical modifications of this group amenable to its photorelease. Therefore, we considered alternatives in other positions of melatonin that could present higher reactivity, as in the case of *N1*. On the one hand, the presence of relatively bulky groups at the *N1*-position decreases the affinity of a ligand for both melatonin receptors, without displaying selectivity towards MT_1_ or MT_2_.^18–20^ On the other hand, indolic nitrogen can be chemically modified with different groups facilitating the synthesis of the target compounds. Therefore, melatonin *N1* was the selected position for the introduction of PPG groups in the structure of these caged derivatives (**Figure 1**).

**Figure 1.**
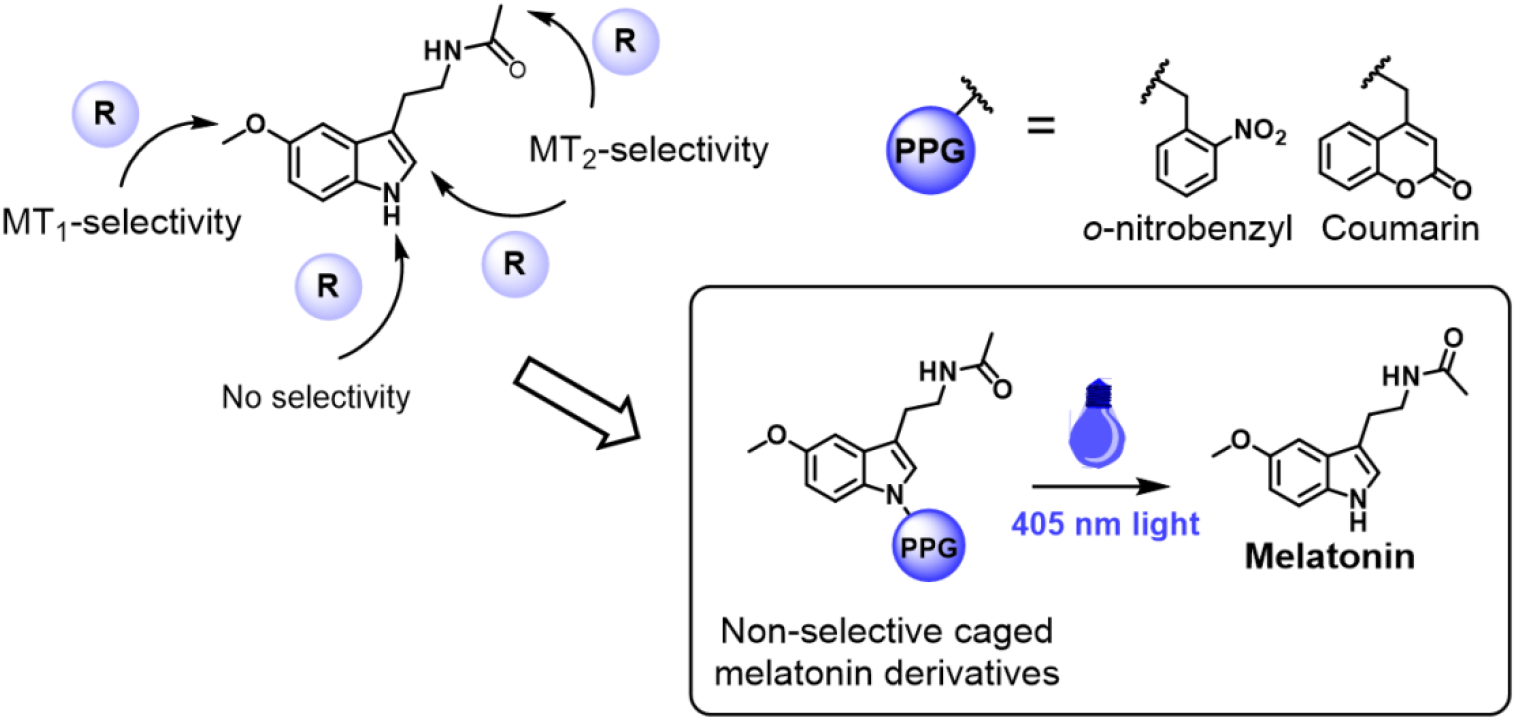
Chemical design of the caged melatonin derivatives presented here. Following the caging strategy, a photoprotective group is attached at the *N1*-position, achieving a decrease on binding affinity for melatonin receptors, without selectivity for MT_1_ or MT_2_. The new light-sensitive molecules release active melatonin upon suitable light conditions.

We aimed to design three compounds (**MCS-0378, MCS-0380, MCS-0381**) containing a carbamate linker between melatonin and the caging group, and one compound (**MCS-0382**) presenting a 4,5-dimethoxy-2-nitrobenzyl (DMNB) group directly attached to melatonin by *N1*-benzylation. Although literature on *N*1-*o*-nitrobenzyl protected indole derivatives is scarce, the contradictory reports in the efficiency of indole photorelease from these precursors^21,22^ prompted us to consider direct alkylation of melatonin indolic nitrogen as a possible alternative to carbamates. Therefore, **MCS-0382** was obtained in low to moderate yields after deprotonation of melatonin with sodium hydride, followed by the rapid addition of DMNB-bromide in DMF at low temperature (**Scheme 2A**). The competitive *N*-benzylation of melatonin acetamide and double alkylation were side-reactions. Two different synthetic pathways were used to obtain the carbamate derivatives, as shown in **Scheme 2B**,**C**. In the first reaction, a two-step, one-pot *N*-acylation of indole was performed, following the methodology described by Macor et al. in 1999.^23^ Accordingly, carbonyldiimidazole (CDI) was used as an activating agent in the presence of a catalytic amount of DMAP. However, this procedure was only valid for the synthesis of **MCS-0378**, which was obtained in 15% overall yield. Poor yields were explained by the main formation of the dimer of melatonin, linked at the *N1*-position by a carbonyl group. In order to improve reaction efficiency, a second synthetic methodology was proposed (**Scheme 2C**). In this reaction, CDI was substituted by 4-nitrophenyl chloroformate in the first step of the synthesis, avoiding the side reaction of melatonin dimerization. The obtained intermediate then reacted with the benzyl alcohol derivative of DMNB or diethylaminocoumarin (DEAC) in ACN, in the presence of catalytic DMAP. These conditions improved slightly the overall yield of the synthesis, and compounds **MCS-0380** and **MCS-0381** were finally obtained in 26 and 53% yields, respectively. The improvement of the synthetic procedures for these compounds was not performed due to the lower photochemical performances of the carbamate linked melatonin derivatives. The subnanomolar activity of melatonin at MT_1_ and MT_2_ receptors required efficient exhaustive purification of the caged derivatives to secure the absence of this bioactive compound in the samples used for biological testing.

Another important aspect to consider is the release of side products upon uncaging.^11^ For instance, photolytic cleavage of the *o-*NB-based PPGs yields originates nitrosobenzene derivatives,^24^ whose possible interference with the biological system must be determined. A similar result is obtained with coumarin-containing PPGs, as they normally generate the corresponding coumarin benzyl alcohol.^25^ In order to test the eventual effect of the cage photolytic products towards melatonin receptors in cells, three light-sensitive phosphate derivatives were synthesized, one for each caging group described herein (*o-*NB, DMNB and DEAC) that release inactive phosphate upon illumination. Details on the synthesis of these molecules are provided on the Supplementary information (**Scheme S2**).

### Photochemical characterization of caged melatonin compounds

UV-Vis absorption spectra was recorded for each compound, in order to determine the interval of wavelengths that promote the uncaging process. Compound **MCS-0378** showed two absorption peaks at λ = 260 nm and λ = 306 nm (**Figure 2A**), which are concordant in agreement with the presence of an *o-*NB moiety. Compound **MCS-0380** displayed an absorption maximum of λ = 380 nm (**Figure 2B**), consistent with the higher absorption wavelengths of coumarin-based PPGs. DMNB-type compounds **MCS-0381** and **MCS-0382** are based on a *o-*NB derivative, but they present two additional methoxy in their structure, which are electron-donating groups (EDGs) that induce a slight bathochromic shift. Therefore, the absorption maxima of these compounds are displaced to longer wavelengths in comparison to **MCS-0378**, with two clear absorption maxima at λ = 306 nm and λ = 350 nm for both cases (**Figures 2C,D**).

**Figure 2.**
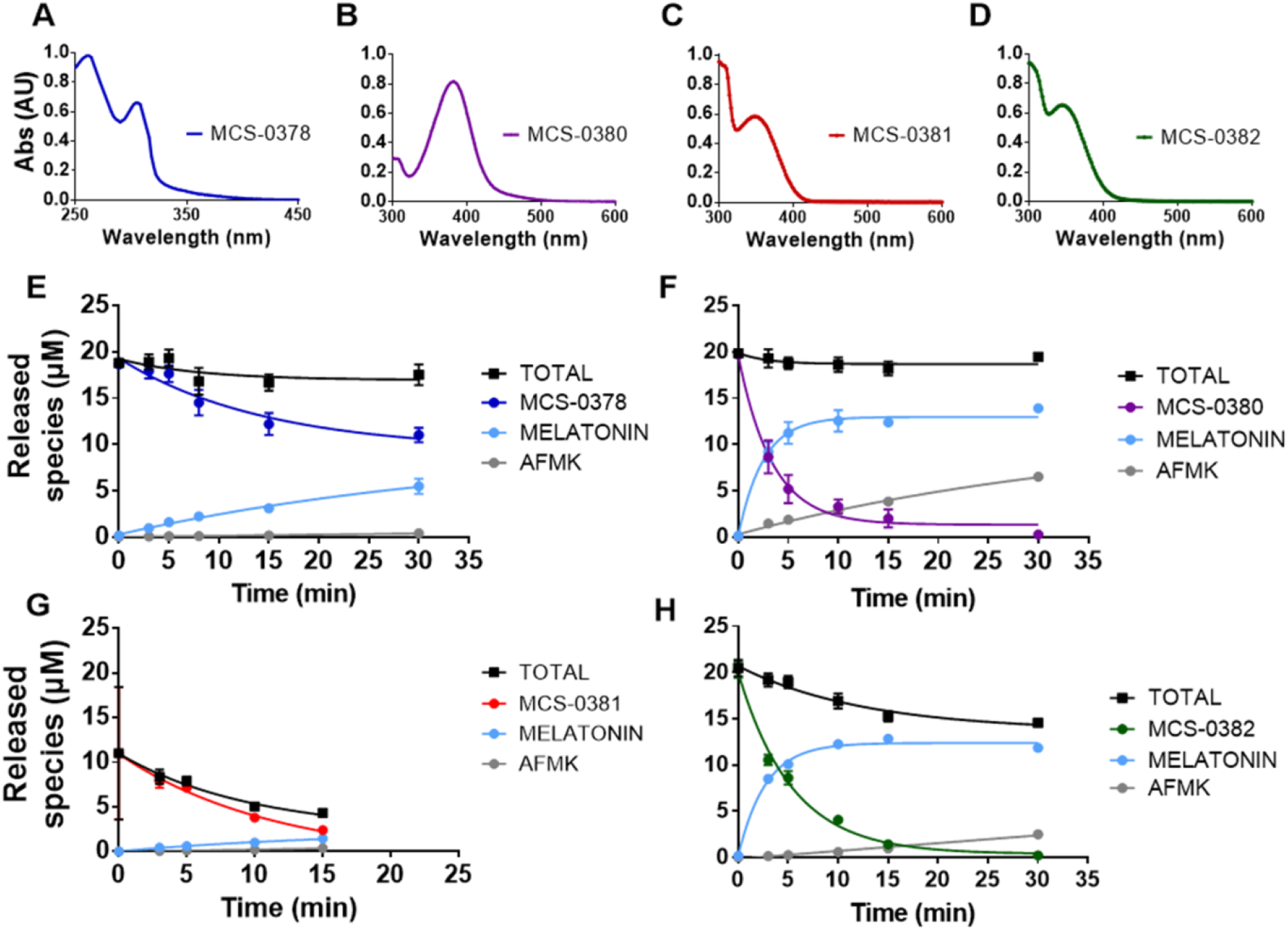
Photochemical characterization of compounds **MCS-0378, MCS-0380, MCS-0381** and **MCS-0382**. UV-Vis absorption spectra of MCS-0378 **(A)**, MCS-0380 **(B)**, MCS-0378 **(C)** and MCS-0382 **(D)**, recorded to a 50 µM solution in 100% DMSO. Photolysis of compounds MCS-0378 **(E)**, MCS-0380 **(F)**, MCS-0378 **(G)** and MCS-0382 **(H)**, performed at λ = 405 nm in a 20 µM solution in PBS:DMSO (98:2). Data are expressed as mean ± SEM of three independent experiments.

Considering the information extracted from the UV-Vis of the different compounds, the photolytic process should be triggered with light of wavelengths ranging from 320 to 400 nm. To avoid the toxicity and cell damage effects associated with UV light, we used a laser with a wavelength in the visible spectrum range (λ = 405 nm, 2.37 mW/mm^2^).

We then monitored photolysis and light stability under conditions similar to those used in biological assays. Aqueous solutions of the caged compounds were illuminated for different periods of time and subsequently analyzed by HPLC-MS. For each time point, we quantified the amount of released melatonin and verified the formation of side products (**Figures 2E-H**). Concentrations of the different species in the sample were calculated with the equation extracted from the calibration curve for each analyte (**Figure S2**).

Compound **MCS-0378** did not present a complete uncaging (**Figure 2E**), as it mainly remained in its caged form even after long light exposures, consistent with their absorption maxima at lower wavelengths. Consequently, the amount of released melatonin at 405 nm was not prominent, reaching concentrations below 5 µM after applying light for 15 min to a 20 µM solution. Interestingly, no photolytic degradation was detected under these conditions.

Compounds **MCS-0380** and **MCS-0382** presented similar uncaging rates and photostability (**Figure 2F,H)**, with a photolytic reaction that was completed after 15 min. In both cases, melatonin was released with relatively high uncaging efficiencies after 5 min irradiation, detecting concentrations between 10 and 13 µM (corresponding to 50% and 65% of the theoretical conversion, respectively). However, melatonin does not entirely account for the phototransformed compound **MCS-0382** (**Figure 2H**). This can be due to the formation of undefined intermediates upon photolysis of *o*-NB-derivatives,^26^ or by the partial reaction of melatonin with the nitrosoaldehyde intermediate, as reported in the *N1*-nitrobenzyl photolytic deprotection of indoles.^21^ Nevertheless, properties of compound **MCS-0382** are remarkable, not only for the high uncaging yields upon light exposure, but also for its chemical stability and improved solubility in aqueous media.

In contrast, the amount of released melatonin from the compound **MCS-0381** was lower than expected (**Figure 2F**), with concentrations that oscillated between 1 and 2 µM after 15 minutes of light exposure. In addition to unfavorable photochemical properties, this compound presented important solubility issues in aqueous solution, requiring high percentages of organic solvent (higher than 20%) to avoid precipitation. Overall, these properties hindered the study of the uncaging process and seriously limited the applicability of **MCS-0381** to further in vitro studies in cells.

Interestingly, apart from the expected release of melatonin we detected the formation of a unique photolysis product, which was identified as *N*^1^-acetyl-*N*^2^-formyl-5-methoxykynuramine (AFMK). AFMK is the main product of the oxidative metabolism of melatonin, and it presents well-established antioxidant properties.^27^ Its formation involves a pyrrole-ring oxidative cleavage of melatonin,^28^ a process that is favored by the action of light.^29^ Therefore, the AFMK concentration was systematically monitored, revealing a minor formation of this photo-oxidation product under our assay conditions (**Figure 2E-H**, grey lines; **Figures S1**).

### Affinity of caged melatonin compounds for MT_1_ and MT_2_ receptors

The affinity of the caged compounds for human melatonin MT_1_ and MT_2_ receptors stably expressed in HEK293 cells was determined in competition binding assays with 2-[^125^I]iodomelatonin (2-[^125^I]-MLT), under dark conditions and after the pre-illumination of the caged compounds for 15 minutes at 405 nm. The reference compound melatonin showed the expected sub-nanomolar affinity for MT_1_ and MT_2_ (pK_i_ = 9.56 ± 0.29 and 9.76 ± 0.45, respectively). All caged compounds presented the expected reduction (>100-fold) in binding affinity, with the exception of compound MCS-0380, which maintained an intermediate affinity for MT_2_ with a pK_i_ of 8.34 ± 0.50 (**Table 1**). The highest shift in affinity (1000-fold) was observed for **MCS-0382** with pK_i_ values in the micromolar range (pK_i_ = 6.32 ± 0.23 and 6.56 ± 0.19 for MT_1_ and MT_2_, respectively).

**Table 1.**
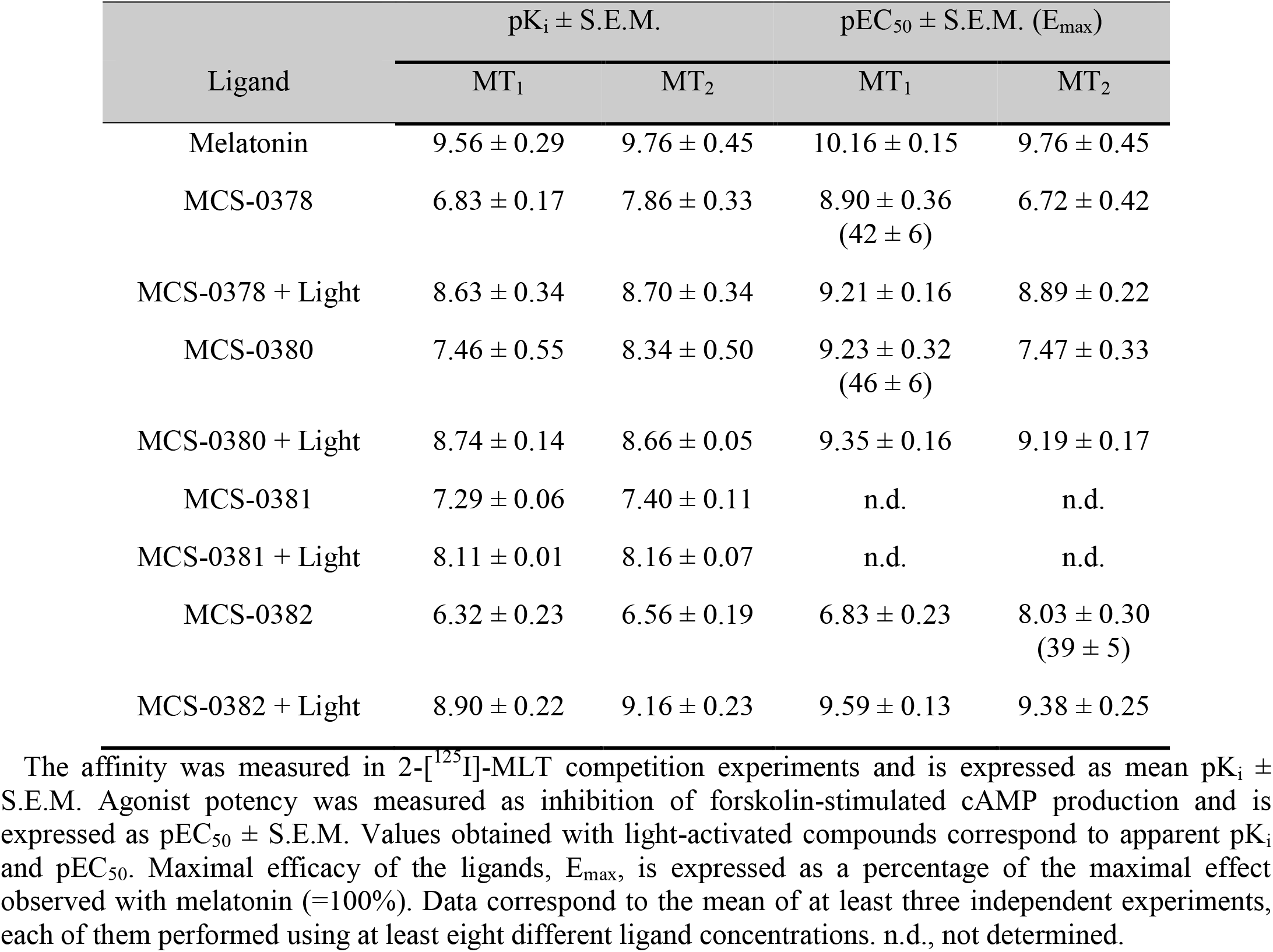
Affinity and agonist potency of MCS-0378, MCS-0380, MCS-0381 and MCS-0382 in HEK293 cells stably expressing MT_1_ and MT_2_.

Upon illumination of the caged compounds for 10 minutes, an increase in apparent affinity was observed on both receptors, confirming the light-mediated generation of biologically active melatonin. Remarkably, melatonin itself was not affected by the light exposure as pK_i_ values remained unchanged (**Table S1**). The increase in the apparent affinity correlates on the uncaging efficiency of each compound and the amount of melatonin generated. Accordingly, compound **MCS**-**0382** presented the most promising properties, with the highest apparent affinity shift from the micromolar range in dark conditions to low nanomolar range upon uncaging (**Figure 3G,H, Table 1**). In contrast, compounds with inferior uncaging efficiencies, like compound **MCS-0381**, did not show an important increase of apparent affinity after uncaging, consistent with the lower amount of released melatonin observed for this compound (**Figure 3E,F, Table 1**). Compounds **MCS-0378** and **MCS-0380** displayed a moderate increase of apparent affinity after light application (**Figure 3A-D**). The narrow window in affinity difference between dark and light conditions of **MCS-0378** and **MCS-0380** is most likely due to the modest loss of affinity by the introduction of the caging groups.

**Figure 3.**
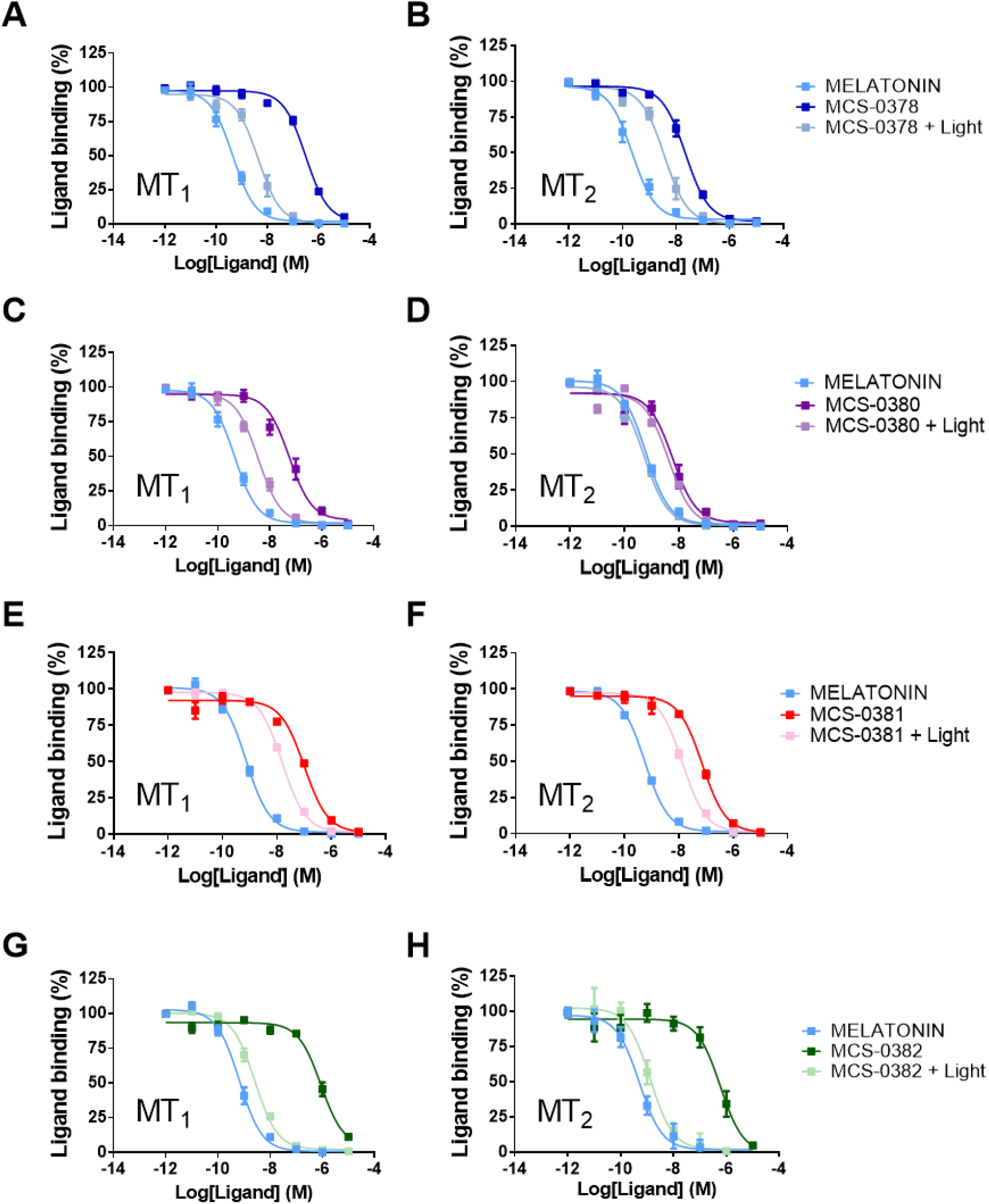
Competition of 2-[^125^I]-MLT binding by melatonin, **MCS-0378, MCS-0380, MCS-0381** and **MCS-0382** before and after light exposure, in crude membranes from HEK293 cells expressing MT_1_ or MT_2_ receptors. Light is applied to the ligands for 10 minutes prior to incubation with the membranes, using the laser as a light source (405 nm, 2.37 mW/mm^2^). Data are expressed as mean ± SEM from 3 to 5 independent experiments (A. Melatonin n = 4, MCS-0378 (Dark) n = 3, MCS-0378 (Light) n = 3; B. Melatonin n = 5, MCS-0378 (Dark) n = 3, MCS-0378 (Light) n = 3; A. Melatonin n = 4, MCS-0378 (Dark) n = 3, MCS-0378 (Light) n = 3; C. Melatonin n = 4, MCS-0380 (Dark) n = 4, MCS-0380 (Light) n = 3; D, E and F: Melatonin, MCS-0380 (Dark), MCS-0380 (Light), MCS-0381 (Dark), MCS-0381 (Light) n = 3; G. Melatonin n = 5, MCS-0382 (Dark) n = 5, MCS-0382 (Light) n = 4; H. Melatonin n = 5, MCS-0382 (Dark) n = 5, MCS-0382 (Light) n = 3). Data are represented as percentage of maximal binding in the absence of compounds and normalized to melatonin maximum effect.

To exclude any impact of other photolytic products generated in the course of the uncaging reaction, we determined the affinity of AFMK and the caging groups for melatonin receptors, before and after light application. Neither the phosphate derivatives of the caging groups (**Figure S4**) nor AFMK (**Figure S5**) showed any significant interference at concentrations up to 1µM. These results exclude any significant impact of these photolytic products on the pharmacological properties of uncaged melatonin on MT_1_ and MT_2_ receptors.

In conclusion, we generated several caged melatonin compounds with variable uncaging efficiencies upon light illumination. The apparent affinity of uncaged melatonin for MT_1_ and MT_2_ receptors is governed by the uncaging efficiency of each compound and no significant interferences of the released cage moiety or photolytic side-products could be detected. The dimethoxynitrobenzyl compound **MCS-0382** shows the most interesting properties with an affinity shift of almost 3 logs upon light illumination on MT_1_ and MT_2_ receptors, useful for biological applications. Its apparent affinity close to pure melatonin is in good accordance to its high uncaging efficiency (65%, 10 min, 405 nm, 2.37 mW/mm^2^) and was selected for cell assays due to the optimal photochemical and receptor affinity properties.

### Functional activity of caged melatonin compounds on MT_1_ and MT_2_ receptors

The well-documented property of melatonin receptor agonists to inhibit intracellular cAMP production was used as a functional readout for our compounds. All caged melatonin compounds presented agonistic properties, i.e. inhibited forskolin-stimulated cAMP production for MT_1_ and MT_2_ receptors prior to light activation (**Table 1**). Whereas pure melatonin showed expected EC_50_ values in the sub-nanomolar range, EC_50_ values for caged compounds were 1-3 logs higher (**Table 1**). **MCS-0378** and **MCS-0380** were partial agonists for MT_1_, and full agonists for MT_2_ (**Figure 4A-D**). In contrast, **MCS-0382** behaved as full agonist for MT_1_ and as partial agonist for MT_2_ (**Figure 4E,F**). Upon light activation at 405 nm, all compounds became full agonists for both receptors in accordance with the generation of melatonin through uncaging. Apparent EC_50_ values correlated with the uncaging efficiency of each compound (**Table 1**). For **MCS-0382**, apparent EC_50_ values were close to that of pure melatonin, in agreement with its superior uncaging efficiency.

**Figure 4.**
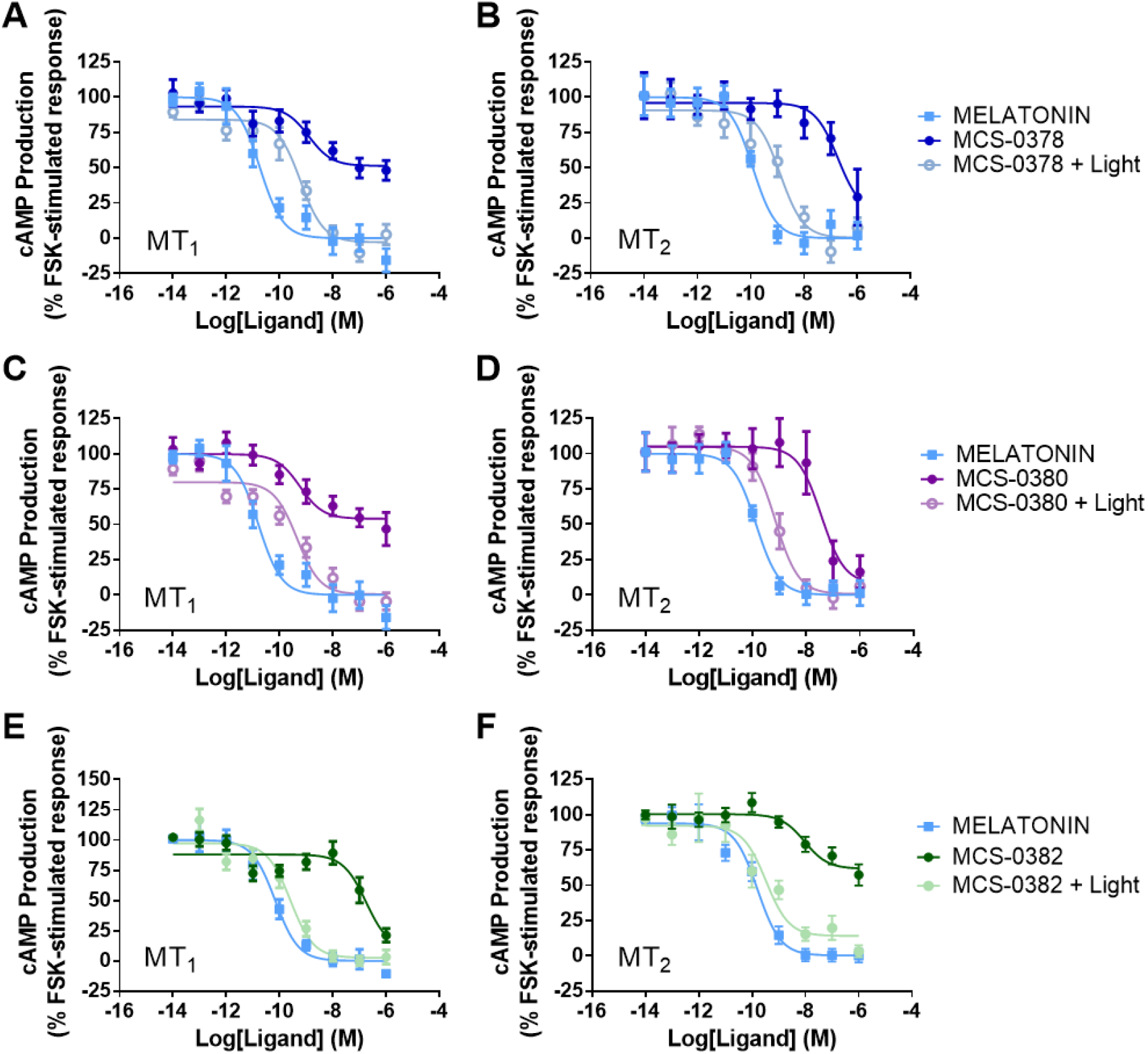
Inhibition of cAMP production by melatonin and compounds **MCS-0378, MCS-0380** and **MCS-0382** before and after light exposure in cells stably expressing MT_1_ (**A**,**C**,**E**) and MT_2_ (**B**,**D**,**F**). Light is applied to the ligands for 10 minutes prior to incubation with the cells, using the laser as a light source (405 nm, 2.37 mW/mm^2^). Data are expressed as mean ± SEM from 4 to 8 independent experiments (**A**. Melatonin n = 7, MCS-0378 (Dark) n = 7, MCS-0378 (Light) n = 7; **B**. Melatonin n = 5, MCS-0378 (Dark) n = 5, MCS-0378 (Light) n = 5; **C**. Melatonin n = 7, MCS-0380 (Dark) n = 7, MCS-0380 (Light) n = 7; **D**. Melatonin n = 5, MCS-0380 (Dark) n = 5, MCS-0380 (Light) n = 5; **E**. Melatonin n = 8, MCS-0382 (Dark) n = 8, MCS-0382 (Light) n = 4; **F**. Melatonin n = 4, MCS-0382 (Dark) n = 4, MCS-0382 (Light) n = 3). Data are presented as percentage of forskolin-stimulated response and normalized to the maximal and minimal melatonin effect. The amplitude of cAMP inhibition before normalization varied between 50 and 75%.

### Electrophysiological analysis of in situ uncaging of MCS-0382 in SCN neurons

We then evaluated the biological activity of the melatonin generated from **MCS-0382** upon light illumination in rat SCN brain slices by monitoring its effect on neuronal excitability in whole-cell recordings. Similar to the reported effect of melatonin in this system at ZT6-11,^30–32^ most SCN neurons (n = 14/21) responded to the application of light-activated **MCS-0382** (**Figure 5**). Six of 21 neurons showed a hyperpolarization response (**Figure 5A**) and 8/21 a depolarization response (**Figure 5B**), while 7/21 remained unresponsive (**Figure 5C**). This heterogeneity resembles that known for pure melatonin and reflects the proportion of melatonin receptor-expressing neurons and the direct and indirect effect of melatonin on neuronal polarization through the modulation of inhibitory GABAergic transmission.^32^

**Figure 5.**
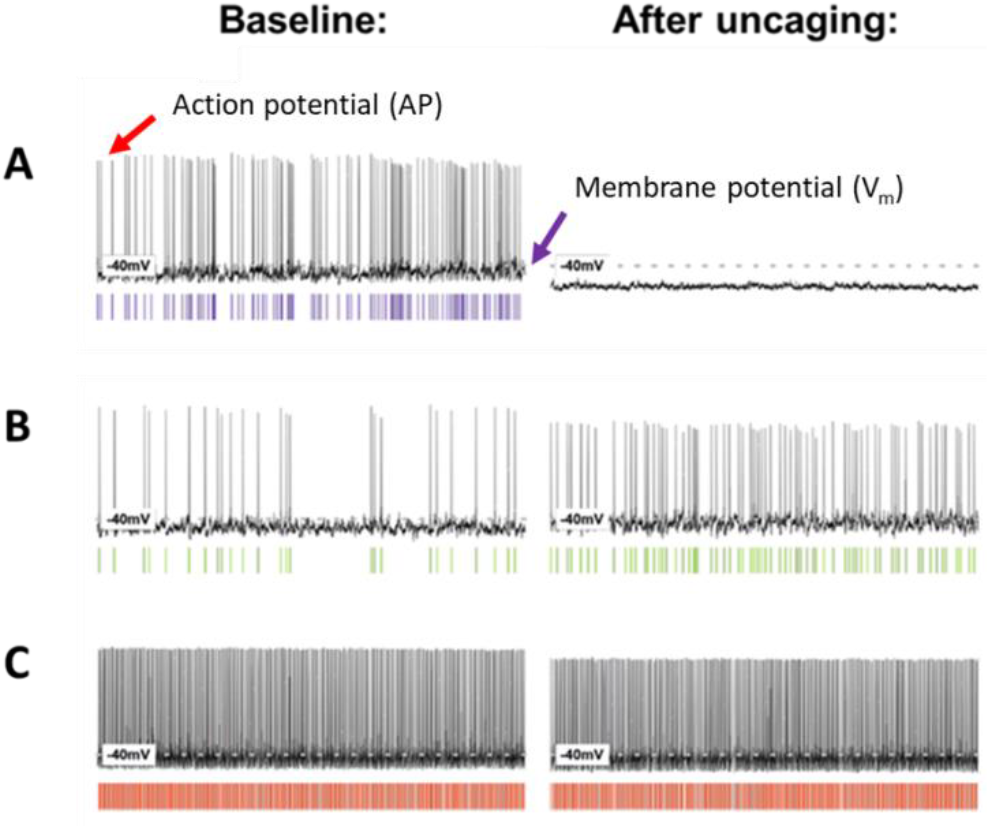
Effects of the application of light on the compound **MCS-0382** on the membrane properties of rat suprachiasmatic nuclei (SCN) neurons. **(A)** Some neurons responded to the light-mediated generation of melatonin with a hyperpolarization (n = 6/21), accompanied by a decrease or elimination of the spontaneous firing rate. **(B)** Other neurons responded to the light-mediated generation of melatonin with a depolarization (n = 8/21), accompanied by an increase of the spontaneous firing rate. **(C)** The remaining neurons did not respond to the light-mediated release of melatonin (n = 7/21). Using the laser (405 nm, 2.37 mW/mm^2^), 1 µM **MCS-0382** was pre-illuminated for 10 minutes prior to application.

In contrast to conventional pharmacology, in photopharmacology, light induces an abrupt change in the concentration of the active molecule in the action site, providing a high spatiotemporal resolution of the process of interest.^33^ Using light to control melatonin receptor activation will be a powerful new pharmacological tool to study the function of these receptors. Light-induced activation of melatonin receptors at specific locations (tissues, brain regions) appears a promising approach to study its effects in experimental biology and pharmacology. Melatonin is known to regulate the sleep/wake cycle and to affect sleep stages but the actual participation of the different brain region(s) expressing melatonin receptors and known to be involved in sleep regulation (i.e., reticular thalamus, the lateral hypothalamus, and the ventrolateral preoptic nucleus) are only poorly defined.^34^ Light activation at specific brain regions followed by electroencephalogram/electromyogram (EEG/EMG) recordings are likely to provide new insights in this respect. A similar approach can be envisioned to better define the brain regions involved in the regulation of pain by melatonin.^35^ Melatonin has been also show to regulate glucose homeostasis but whether this occurs through peripheral or central effects remain poorly known.^36^ Indeed, melatonin is known to regulate the circadian rhythm of the biological master clock located in the hypothalamic suprachiasmatic nucleus and to affect glucose homeostasis by directly acting on glucose sensitive peripheral organs such as the liver or the pancreas. Local activation of melatonin receptors is likely to detangle the respective contributions of these organs.

Our caged melatonin compounds present not only a proof-of-concept for the feasibility of melatonin as a photo-pharmacological tool but also for the development of other photo-activable molecules and sub-type selective melatonin receptors ligands. For instance, melatonin derivatives containing caging groups that undergo photolysis at longer and even less invasive wavelengths can be envisioned, such as BODIPY, or the melatonin moiety replaced by an antagonistic or receptor type-selective molecule.^10^ Caged MT_1_- and MT_2_-selective molecules would be of particular interest to further decipher the specific and opposing effects of the two melatonin receptors on sleep.^37^ A further development of our caged melatonin compounds could be the attachment of moieties that target specific subcellular compartments. The latter provides a higher degree of optical control, and it has been previously applied to release caged lipids from organelles such as mitochondria, lysosomes or endoplasmic reticulum.^38,39^ Subcellular compartment-targeted molecules might be of particular interest to study the role of the previously described mitochondrial MT_1_ receptors.^40,41^

## CONCLUSION

In this work, we describe the synthesis and functional characterization of the first family of photo-activable melatonin receptor ligands. Following the caging strategy, three different PPGs have been selected to generate compounds **MCS-0378, MCS-0380, MCS-0381** and **MCS-0382**, by attachment of the caging group at the *N1*-position of melatonin. With the exception of **MCS-0381**, all compounds release active melatonin with relatively high yields, which is reflected by the excellent apparent EC_50_ values for the inhibition of cAMP production of the compounds upon 405 nm illumination. The presence of the caging groups induced an important decrease in binding affinity of caged melatonin compounds for both receptors giving 2-3 log concentration windows that are useful for light experiments in cells and tissues. These results reveal the interest of *N1*-substitutions for the development of non-selective low-affinity ligands for melatonin receptors. **MCS-0382** showed the most interesting properties as it is chemically stable, highly soluble in aqueous media, and displays an excellent uncaging efficiency and a more than 100-fold difference in affinity before and after photolysis. The biological activity of melatonin generated through light-induced uncaging was demonstrated in two cellular assays (ligand binding and signaling assays) and in rat SCN brain slices. Collectively, these results validate the caged melatonin derivative **MCS-0382** as photopharmacological tool for cellular and physiological studies.

## EXPERIMENTAL SECTION

### Organic synthesis

All chemicals and solvents were obtained from commercial sources and used without purification, except anhydrous solvents, which were previously treated with a system of solvent purification (*PureSolvEN*^™^), degassed by purging with inert gases and dried over molecular sieves. Reactions were monitored by thin layer chromatography (TLC) on silica gel (60 F, 0.2 mm, ALUGRAM Sil G/UV_254_ *Macherey-Nagel*) by visualization under 254 and/or 365 nm lamp. Compounds without chromophores were visualized using ethanolic solution of phosphomolybdic acid by heating. Alternatively, Nuclear Magnetic Resonance (NMR) was also used as a technique for reaction monitoring. When purification was required, *flash* column chromatography was performed on silica-gel 60 (Panreac, 40-63 microns RE). Reversed-phase column *flash* chromatography was done on silica-gel C18 (SNAP KP-C18-HS, 50μ, *Biotage*) and automated with *Isolera*^*™*^ *One* with UV-Vis detection (*Biotage*).

Compound characterization by NMR spectroscopy was performed with *Variant-Mercury 400 MHz* spectrometer. Chemical shifts d are reported in parts per million (ppm) from tetramethylsilane as internal standard and using residual non-deuterated solvent signal as reference (Chloroform-*d* d = 7.26 ppm (^1^H), d = 77.16 ppm (^13^C); Methanol-*d*_*4*_ d = 4.87 ppm, d = 3.31 ppm (^1^H), d = 49.3 ppm (^13^C)). The following abbreviations were used to designate multiplicities: s=singlet, d=doublet, t=triplet, q=quartet, m=multiplet, q=quintuplet, br=broad, dd=double-doublet, ddd=double-double-doublet, dt=double-triplet, td=triple-doublet. Coupling constants (*J*) were expressed in Hz. Signals were assigned as far as possible by means of two-dimensional NMR spectroscopy: 1H-1H-COSY, HSQC and HMBC.

Purity of compounds were determined by HPLC using two different methods (Method A and Method B). In both cases, purity is given as % of absorbance at 254 nm.

Method A uses a *Dionex Ultimate 3000SD* HPLC (*Thermo Fischer Scientific*), which is coupled to a PDA detector and to a mass spectrometer *LTQ XL ESI-ion trap* (*Thermo Fischer Scientific*). For this method, column used was *ZORBAX Extend C18* (2.1 × 50 mm, 3.5 µm; P.N. 735700-902). The flow rate was 0.9 mL/min, column temperature was fixed to 30 °C and total runtime is 10 min. The mobile phase used was a mixture of A = formic acid 0.05% in water and B = formic acid 0.05% in acetonitrile (ACN), with the method described as follows: from 5% of B to 90%B in 5 min, 90% of B for 2 min, from 90% of B to 100% of B in 1 min and 100% of B for 2 min. UV-Vis spectra were collected every 0.2 s between 190 and 800 nm and bands are % of maximal absorbance. Data from mass spectra were analyzed by electrospray ionization in positive and negative mode between 50 and 2000 Da, using Xcalibur software version 2.2 (*Thermo Fischer Scientific*). Method B uses a *Waters2795 Alliance* HPLC, coupled to a DAD detector (*Agilent 1100*) and to an ESI *Quattro micro* MS detector (*Waters*). For this method, column used was *ZORBAX Eclipse Plus C18* (4.6 × 150 mm, 3.5 µm). The flow rate was 0.5 mL/min, column temperature was fixed to 35 °C and total runtime is 10 min. The mobile phase used was a mixture of A = formic acid 0.05% in water and B = formic acid 0.05% in ACN, with the method described as follows: 5% of B for 0.5 min, from 5% B to 100% B in 5 min, 100% of B for 1.5 min, from 100% of B to 5% of B in 1 min, and 5% of B for 2 min. UV-Vis spectra were collected every 0.2 s between 210 and 600 nm and bands are % of maximal absorbance. Data from mass spectra were analyzed by electrospray ionization in positive and negative mode every 0.3 s between 150 and 1500 Da, using MassLynx software version 4.1 (*Waters*).

High-resolution mass spectra (HRMS) and elemental composition were analyzed by FIA (flux injected analysis) using Ultrahigh-Performance Liquid Chromatography (UPLC) *Aquity (Waters)* coupled to LCT Premier Orthogonal Accelerated Time of Flight Mass Spectrometer (TOF) (*Waters*). Data from mass spectra were analyzed by electrospray ionization in positive and negative mode using MassLynx software version 4.1 (*Waters*). Spectra were scanned between 50 and 1500 Da with values every 0.2 seconds and peaks are given m/z. Melting points were taken on open capillary tubes and measured with Melting Point B-545 (*Büchi*), ramp 0.5 °C/min with a digital temperature measurement. IR spectra were registered in chloroform solution and recorded using *Thermo Nicolet Avatar 360 FT-IR* Spectrometer.

2-nitrobenzyl 3-(2-acetamidoethyl)-5-methoxy-*1H*-indole-1-carboxylate (**MCS-0378**). To a solution of commercially available melatonin (90 mg, 0.38 mmol, 1 eq) in dry ACN (0.2 M) under an inert atmosphere, a solution of di(1H-imidazol-1-yl)methanone (70.4 mg, 0.42 mmol, 1.1 eq) and *N,N*-dimethylpyridin-4-amine (11.6 mg, 0.09 mmol, 0.25 eq) in dry ACN (4.4 mL) were added dropwise at room temperature. The mixture was stirred for 30 minutes at 40 °C, and a solution of 2-nitrobenzyl alcohol (83 mg, 0.53 mmol, 1.4 eq) in dry ACN (0.5 mL, 1 M) was then added. The resulting solution was heated under reflux with magnetic stirring for two days. The reaction was then completed, quenched with HCl 1 M (30 mL) and the aqueous layer was extracted with EtOAc (4 × 25 mL). Combined organic layers were washed with brine (2 × 40 mL), dried over anhydrous Na_2_SO_4_, filtered and evaporated under reduced pressure. The crude was first purified by column chromatography through silica gel using DCM/MeOH (98:2), in order to discard the benzyl bromide derivatives and melatonin. The product-containing fractions were re-purified by column chromatography through silica gel using DCM/MeOH (97:3), giving the expected product as a white solid (22.3 mg, 15%).

^1^H NMR (400 MHz, Chloroform-*d*): d = 8.15 (d, *J* = 8.2 Hz, 1H), 8.04 (s, 1H), 7.71 – 7.67 (m, 2H), 7.55 (ddd, *J* = 8.6, 5.5, 3.3 Hz, 1H), 7.42 (s, 1H), 7.01 (d, *J* = 2.5 Hz, 1H), 6.96 (dd, *J* = 8.9, 2.5 Hz, 1H), 5.83 (s, 2H), 5.64 (s, 1H), 3.87 (s, 3H), 3.59 (q, *J* = 6.5 Hz, 2H), 2.89 (t, *J* = 6.8 Hz, 2H), 1.97 (s, 3H). ^13^C NMR (101 MHz, Chloroform-*d*): d = 170.38 (2C), 156.53 (2C), 134.09, 131.48 (2C), 129.57, 129.50 (2C), 125.41, 119.26 (2C), 116.23, 113.71, 102.14, 65.24, 55.93, 39.08, 25.31, 23.52. HPLC-PDA-MS (using method A): RT = 2.66 min, *λ*_max_ = 209, 243, 299 nm; purity 98% (254 nm). HRMS calculated for C_21_H_22_N_3_O_6_: 412.1509 [M+1]^+^, found: 412.1489; mp 133.2-133.8°C

4-nitrophenyl 3-(2-acetamidoethyl)-5-methoxy-*1H*-indole-1-carboxylate (**1**). To a solution of commercially available melatonin (96 mg, 0.40 mmol) in dry ACN (1.5 mL, 0.3 M) under an inert atmosphere, a solution of 4-nitrophenyl carbonochloridate (105 mg, 0.50 mmol) in dry ACN (0.5 mL, 1 M), a solution of DMAP (12.34 mg, 0.10 mmol) in dry ACN (0.2 mL, 0.5 M) and triethylamine (70 µL, 0.50 mmol) were added dropwise. The resulting mixture was heated under reflux overnight with magnetic stirring. The reaction was then quenched with HCl 1 M (50 mL) and the aqueous layer was extracted with EtOAc (4 × 25 mL). Combined organic layers were washed with brine (2 × 40 mL), dried over anhydrous Na_2_SO_4_, filtered and evaporated under reduced pressure. The crude was purified by column chromatography through silica gel using DCM/MeOH (98:2), giving the expected product as beige crystals (78 mg, 49%).

^1^H-NMR (400 MHz, Chloroform-*d*): d = 8.39 – 8.30 (m, 2H), 8.08 (s, 1H), 7.53 (s, 1H), 7.52 – 7.46 (m, 2H), 7.07 (d, *J* = 2.4 Hz, 1H), 6.99 (dd, *J* = 9.0, 2.5 Hz, 1H), 5.70 (s, 1H), 3.89 (s, 3H), 3.62 (q, *J* = 6.7 Hz, 2H), 2.93 (t, *J* = 7.0 Hz, 2H), 1.98 (s, 3H). ^13^C NMR (101 MHz, Chloroform-*d*): d = 170.37, 156.94, 154.94, 145.80, 131.73, 130.13, 125.55 (2C), 122.81, 122.50 (3C), 120.43, 116.33, 113.92, 102.50, 55.92, 39.12, 25.41, 23.51. HPLC-PDA-MS (using method B): RT = 3.55 min, *λ*_max_ = 210, 244, 263, 300 nm; purity 95% (254 nm). HRMS calculated for C_20_H_20_N_3_O_6_: 398.1352 [M+1]^+^, found: 398.1356; mp 182.1-182.2 °C.

(7-(diethylamino)-2-oxo-2H-chromen-4-yl)methyl -(2-acetamidoethyl)-5-methoxy-*1H*-indole-1-carboxylate (**MCS-0380**). To a stirred solution of 4-nitrophenyl 3-(2-acetamidoethyl)-5-methoxy-*1H*-indole-1-carboxylate **1** (52 mg, 0.13 mmol, 1 eq) in dry ACN (1 mL) under an argon atmosphere, a solution of DEAC-OH (38 mg, 0.14 mmol, 1.3 eq) and *N,N*-dimethylpyridin-4-amine (4.01 mg, 0.03 mmol, 0.25 eq) in dry ACN (0.5 mL) were added. The mixture was immediately protected from light and heated under reflux under magnetic for three days. Then, the reaction was quenched with a HCl 1M solution (50 mL), and the aqueous layer was extracted with EtOAc (4 × 25 mL). Combined organic layers were over anhydrous Na_2_SO_4_, filtered and evaporated under reduced pressure. The crude was first purified by column chromatography through silica gel using DCM/MeOH (97:3), in order to discard the benzyl derivatives and melatonin. The product-containing fractions were re-purified using the same conditions as before, DCM/MeOH (98:2), giving the expected product as a yellow solid (35 mg, 53%).

^1^H-NMR (400 MHz, Chloroform-*d*): d = 8.06 (s, 1H), 7.44 (s, 1H), 7.37 (d, J = 8.9 Hz, 1H), 7.04 (d, J = 2.5 Hz, 1H), 6.96 (dd, J = 9.0, 2.5 Hz, 1H), 6.62 (dd, J = 9.0, 2.6 Hz, 1H), 6.54 (d, J = 2.6 Hz, 1H), 6.20 (s, 1H), 5.62 (s, 1H), 5.53 (s, 2H), 3.87 (s, 3H), 3.59 (q, J = 6.6 Hz, 2H), 3.43 (q, J = 7.1 Hz, 4H), 2.90 (t, J = 6.9 Hz, 2H), 1.97 (s, 3H), 1.21 (d, J = 7.0 Hz, 6H). ^13^C NMR (101 MHz, Chloroform-*d*): d = 170.34, 161.82, 156.61, 156.48, 150.71, 148.92, 131.47, 124.55, 122.88, 119.65 (2C), 116.19 (2C), 113.83, 109.18, 107.00, 105.95, 102.22, 98.27, 63.67, 55.92, 45.09 (2C), 39.23, 25.34, 23.54, 12.54 (2C). HPLC-PDA-MS (using method B): RT = 3.83 min, *λ*_max_ = 210, 244, 386 nm; purity > 98% (254 nm). HRMS calculated for C_28_H_32_N_3_O_6_: 506.2291 [M+1]^+^, found: 506.2291; mp 173.1-173.6 °C.

4,5-dimethoxy-2-nitrobenzyl 3-(2-acetamidoethyl)-5-methoxy-*1H*-indole-1-carboxylate (**MCS-0381**). To a stirred solution of 4-nitrophenyl 3-(2-acetamidoethyl)-5-methoxy-*1H*-indole-1-carboxylate **1** obtained in the first step (132 mg, 0.30 mmol, 1 eq) in dry ACN (3 mL) under an argon atmosphere, a solution of 4,5-dimethoxy-2-nitrobenzyl alcohol (99 mg, 0.45 mmol, 1.3 eq) and *N,N*-dimethylpyridin-4-amine (9.26 mg, 0.08 mmol, 0.25 eq) in dry ACN (1 mL) were added. The mixture was immediately protected from light and heated under reflux under magnetic stirring for six days. Then, the reaction was quenched with a HCl 1M solution (50 mL), and the aqueous layer was extracted with EtOAc (4 × 25 mL). Combined organic layers were over anhydrous Na_2_SO_4_, filtered and evaporated under reduced pressure. The crude was first purified by column chromatography through silica gel using DCM/EtOAc (95:5), in order to discard the benzyl derivatives and melatonin. The product-containing fractions were re-purified using the same conditions as before: DCM/MeOH (98:2), giving the expected product as a pale-brown solid (37.5 mg, 26%).

^1^H-NMR (400 MHz, Chloroform-*d*): d = 8.04 (s, 1H), 7.75 (s, 1H), 7.42 (s, 1H), 7.10 (s, 1H), 7.01 (d, *J* = 2.0 Hz, 1H), 6.95 (dd, *J* = 9.0, 2.5 Hz, 1H), 5.82 (s, 2H), 5.60 (s, 1H), 3.97 (s, 3H), 3.95 (s, 3H), 3.87 (s, 3H), 3.58 (q, *J* = 6.6 Hz, 2H), 2.88 (t, *J* = 6.9 Hz, 2H), 1.96 (s, 3H). ^13^C NMR (101 MHz, Chloroform-*d*): d = 170.37, 156.50, 153.46, 148.93, 140.39, 131.36, 130.13, 125.93, 123.05, 122.85, 119.19, 116.19, 113.71, 111.59, 108.60, 102.12, 65.63, 56.70, 56.63, 55.92, 39.14, 25.32, 23.49. HPLC-PDA-MS (using method A): RT = 2.60 min, *λ*_max_ = 195, 244, 301, 344 nm; purity 97% (254 nm). HRMS calculated for C_23_H_26_N_3_O_8_: 472.1720 [M+1]^+^, found: 472.1736; mp 205.7-206.3 °C.

*N*-(2-(1-(4,5-dimethoxy-2-nitrobenzyl)-5-methoxy-*1H*-indol-3-yl)ethyl)acetamide (**MCS-0382**). To a suspension of sodium hydride (17.82 mg, 0.45 mmol, 1.1 eq) in dry DMF (1 mL, 0.45 M) under an argon atmosphere at 0 °C, a solution of commercially available melatonin (97 mg, 0.41 mmol, 1 eq) in dry DMF (0.6 mL, 0.7 M) was added dropwise. The ice/water bath was removed, and the reaction was stirred for 2 hours while the temperature increased to room temperature. Then, an excess of dry DMF (3.5 mL) was added, and the reaction was cooled to -40 °C with an acetone/dry ice bath. A solution of 1-(bromomethyl)-4,5-dimethoxy-2-nitrobenzene (150 mg, 0.53 mmol, 1.3 eq) in dry DMF (0.9 mL, 0.6 M) was added dropwise into the brown mixture, and it was immediately protected from light. The reaction was stirred overnight without exceeding a temperature of 10 °C. The reaction was then quenched with water (25 mL), and the aqueous layer was extracted with EtOAc (3 × 25 mL). The organic layers were washed with brine (4 × 30 mL), dried over anhydrous MgSO_4_, filtered and evaporated under reduced pressure. The crude was first purified by column chromatography through silica gel using DCM/MeOH (95:5), in order to discard the benzyl derivatives and melatonin. The product-containing fractions were re-purified using DCM/EtOAc/MeOH (96:1:3) as eluents, giving the expected product as a pale-brown solid (61.3 mg, 35%).

^1^H-NMR (400 MHz, Chloroform-*d*): d = 7.73 (s, 1H), 7.07 (d, *J* = 2.4 Hz, 1H), 7.03 (d, *J* = 8.9 Hz, 1H), 6.95 (s, 1H), 6.84 (dd, *J* = 8.9, 2.4 Hz, 1H), 5.80 (s, 1H), 5.70 (s, 1H), 5.67 (s, 2H), 3.93 (s, 3H), 3.86 (s, 3H), 3.58 (q, *J* = 6.6 Hz, 2H), 3.44 (s, 3H), 2.96 (t, *J* = 6.9 Hz, 2H), 1.96 (s, 3H). ^13^C NMR (101 MHz, Chloroform-*d*): d = 170.26, 154.39, 154.06, 148.10, 139.51, 132.22, 129.57, 128.57, 127.05, 112.78, 112.75, 110.72, 109.70, 108.41, 101.04, 56.54, 56.19, 56.01, 48.20, 40.12, 25.49, 23.50. HPLC-PDA-MS (using method A): RT = 2.53 min, *λ*_max_ = 235, 282, 303, 344 nm; purity 98% (254 nm). HRMS calculated for C_22_H_26_N_3_O_6_: 428.1822 [M+1]^+^, found: 428.1823; mp 131.3-131.4 °C.

### Photochemistry

UV-Vis absorption spectra of a 100 µM solution in DMSO of each compound were recorded using the Infinite M1000 Tecan microplate reader (λ = 250 – 800 nm). Irradiation experiments to trigger uncaging were performed in a 96-well white plate using a *BlueClassic* laser (TorLaser, Spain) to irradiate the samples from top (λ = 405 nm, 2.37 mW/mm^2^). All samples were prepared with a concentration of 20 µM in aqueous buffer, containing from 2 to 10% DMSO depending on their solubility. 200 µL of these solutions were irradiated for different periods of time (t = 0, 3, 5, 8, 10, 15 and 30 min) and then analyzed by HPLC-MS to monitor photolysis. These analyses were performed in a *Dionex Ultimate 3000SD* HPLC (*Thermo Fischer Scientific*), which is coupled to a PDA detector and to a mass spectrometer *LTQ XL ESI-ion trap* (*Thermo Fischer Scientific*). Conditions of the analysis are those for Method A (see Organic Chemistry section). Calibration curves of all quantifiable species were determined in each experiment, by analyzing a minimum of seven dilutions of each compound, prepared at different concentrations ranging from 0.2 to 50 µM. Curves were fit by plotting the peak area of the analyte versus the concentrations of the analyte with least-squares linear regression. All experiments were performed at least in triplicate.

### Illumination experiments

In order to study the binding affinity and functional properties under light conditions, solutions of the compounds were prepared in aqueous buffer, with concentrations ranging from 10 to 30 µM. 250 µL of solution were deposited in a 96-well white plate and light was applied from top of each well individually, using the same laser source as for the uncaging experiments (see above). Immediately after illumination, the compound was added to the cell system. All the experiments were performed in the dark.

### Cell culture

HEK293 cells were grown in complete medium (Dulbecco’s modified Eagle’s medium supplemented with 10% (v/v) fetal bovine serum (FBS), 4.5g/L glucose, 100 U/mL penicillin, 0.1mg/mL streptomycin and 1mM glutamine) (Invitrogen, CA, USA). Cells were maintained at 37 °C (95% O_2_, 5% CO_2_).

### Crude membrane preparation

Crude membranes were prepared as previously described.^42–44^ The expression of MT_1_ and MT_2_ receptors in the crude membranes was quantified by BCA assay, using Pierce^™^ BCA Protein Assay kit (ThermoFischer Scientific, Whaltham, MA, USA) and following the manufacturer’s instructions.

### Radioligand binding experiments

Radioligand binding assays were performed in 75 mM Tris (pH 7.4), 12 mM MgCl_2_, 5 mM EDTA and 2-[^125^I]-MLT as radioligand (PerkinElmer, Waltham, MA, USA), using membranes extracted from HEK293 cells stably expressing human MT_1_ or MT_2_ receptors. Saturation binding experiments were performed in the range of 1-1000 pM, and specific binding was defined as binding displaced by 10 µM MLT. Competition curves were performed by simultaneous incubation of 200 pm 2-[^125^I]-MLT and increasing concentrations of the respective ligands. Assays were carried out in duplicates for 120 at 37 °C, followed by a rapid filtration through glass fiber filters (Whatman, Clifton, NJ, USA). Filter-retained radioactivity was determined with a γ-counter LB2111 (Berthold Technologies, Bad Wildbad, Germany). Competition curves were fitted with a one-site non-linear regression to determine IC_50_ values, using GraphPad Prism software version 6.0 (San Diego, CA, USA). Data were represented as percentage of maximal binding in the absence of compounds and normalized to melatonin maximum effect. K_i_ values were calculated from IC_50_ values using the Cheng-Prussof formula: K_i_ = IC_50_/[1 + (L/K_d_)], where L represents the 2-[^125^I]-MLT concentration and K_d_ the dissociation constant obtained in the corresponding radioligand saturation assays. K_d_ values were 198 ± 30 and 211 ± 19 pM for MT_1_ and MT_2_, respectively, obtained from three independent saturation binding experiments.

### Accumulative cAMP assay

The accumulative cAMP assay was performed as previously described,^45^ using the CisBio cAMP-G_i_ kit (Cisbio Bioasays, Codolet, France). Briefly, HEK293 cells stably expressing MT_1_ or MT_2_ receptors were dispensed into a 384-well plate (5000 cells per well) and stimulated with 1 µM forskolin in the presence of increasing concentrations of melatonin or the ligands of interest, in PBS buffer supplemented with 1 mM IBMX (Sigma-Aldrich, St Quentin, France) for 30 min at room temperature. Cells were then lysed for 1 h at room temperature, and cAMP levels were determined following the manufacturer’s instructions. The plate was read using the Infinite F500 Tecan microplate reader. Data were fitted by non-linear regression to determine E_max_ and EC_50_ values and normalized to forskolin-induced response (100%) using GraphPad Prism software.

### Animals

Male Sprague-Dawley rats (Charles River, UK), 28 to 35 days old, were housed with free access to standard chow and water in a temperature-controlled environment under 12/12h light/dark conditions with lights on at 8 A.M.. All procedures were performed in accordance with the U.K. Animals (Scientific Procedures) Act 1986 and local ethical approvals.

### Slice preparation

For electrophysiological experiments rats (n=5) were euthanized with sodium pentobarbital and transcardially perfused with ice-cold and oxygenated (95%O2/5%CO_2_) ‘slicing’ solution containing (in mM) sucrose (214), KCl (2.0), NaH_2_PO4 (1.2), NaHCO_3_ (26), MgSO_4_ (1.3), CaCl_2_ (2.4), D-glucose (10). Following decapitation, the brain was extracted and the meninges gently removed. The brain was blocked and glued to a vibratome (Campden Instruments, Loughborough, Leics., UK) where 250 μm thick coronal slices of the hypothalamus containing the SCN were prepared. Slices were immediately transferred to “recording” artificial cerebrospinal fluid (aCSF) containing (in mM) NaCl (127), KCl (2.0), NaH_2_PO_4_ (1.2), NaHCO_3_ (26), MgCl_2_ (1.3), CaCl_2_ (2.4), D-glucose (10), in a continuously oxygenated holding chamber at 35°C for a period of 25 min. Subsequently, slices were allowed to recover in ‘recording’ solution at room temperature for a minimum of 1h before recording.

### Whole-cell patch clamp recordings

For whole-cell recordings, slices were transferred to a submerged chamber and placed on an elevated grid that allows perfusion both above and below the slice. An Olympus BX-51 WI upright microscope (Olympus, Southend-on-Sea, Essex, UK) was used for infrared - differential interference contrast visualization of cells.

Recordings were performed at room temperature (22°C) and slices were continuously perfused with oxygenated ‘recording’ solution (as above) at a rate of ca. 5 ml/min, unless otherwise described. All pharmacological compounds were dissolved in ‘recording’ solution and bath applied. Recording solution containing 1μM MCS-0382 was exposed to blue light (405 nm, 2.37 mW/mm^2^) for 10 minutes prior to application.

Whole cell current- and voltage-clamp recordings were performed with pipettes (3-7MΩ when filled with intracellular solution) made from borosilicate glass capillaries (World Precision Instruments, Hitchin, Herts, UK) pulled on a P-97 Flaming/Brown micropipette puller (Sutter, Novato, CA, USA). The intracellular recording solution contained (in mM) K-gluconate (140), KCl (10), HEPES (10), EGTA (1), Na_2_ATP (2), pH 7.3 (with KOH). Recordings were performed using a Multiclamp 700B amplifier and pClamp11 software (Molecular Devices, San Jose, CA, USA). Slow and fast capacitative components were automatically compensated for. Access resistance was monitored throughout the experiments, and neurons in which the series resistance was >25 MΩ or changed >15% were excluded from the statistics. Liquid junction potential was 16.4mV and not compensated. The recorded current was sampled at 10 kHz and filtered at 2 kHz unless otherwise stated.

### Data and statistical analysis

The data and statistical analysis comply with the recommendations on experimental design and analysis in pharmacology,^46^ using GraphPad Prism software. Data are presented as means ± SEM of *n* independent experiments, performed at least in duplicates to ensure the reliability of single values. IC_50_, EC_50_ and E_max_ values were obtained following non-linear regression (curve fit) with four parameters of data from a minimum of eight different concentrations per experiment, repeated at least three times independently. All assays performed here were previously validated and demonstrated the robustness and variability of the procedure using this number of independent experiments for concentration–response curves. When possible, data were normalized to maximal and minimum responses using melatonin response as a reference in order to avoid unwanted sources of variations, as differences in the amplitude of melatonin effect between independent experiments.

## Supporting information

Additional synthetic procedures, photochemical and pharmacological data

## ASSOCIATED DATA

### Supporting information

Additional Photochemical and Pharmacological data and Compound Characterization Spectra (NMR and HPLC-MS) (PDF)

Molecular Formula Strings (CSV)

## AUTHOR INFORMATION

### Author Contributions

All authors have given approval to the final version of the manuscript. Conceptualization, A.L., and R.J.; Chemical synthesis, G.S.B., C.S.; Biochemical investigations, G.S.B., D.L.; Supervision, H.P., A.L., R.J.; Writing – Original Draft, Review and Editing, G.S.B., A.L., and R.J.; Funding Acquisition, R.J, A.L. D.L. and H.D.P. designed the electrophysiological studies and D.L. made the recordings and analyzed the whole-cell recordings.

### Notes

The authors declare no competing financial interest.

## ACKNOWLEDGEMENTS

This work was supported by Agence Nationale de la Recherche (ANR-19-CE16-0025-01 " mitoGPCR " to R.J.), Institut National de la Santé et de la Recherche Médicale (INSERM), Centre National de la Recherche Scientifique (CNRS) and by Ministerio de Ciencia e Innovación (MCIN), Agencia Estatal de Investigación (AEI) and ERDF-FEDER European Fund (projects CTQ2017-89222-R and PID2020-120499RB-I00) funded by MCIN/AEI/ 10.13039/501100011033 and by “ERDF A way of making Europe” by the European Union, and by the Catalan government (2017SGR1064). R.J. was supported by the Fondation de la Recherche Médicale (Equipe FRM DEQ20130326503), and ANR-21-CE18-00XX " alloGLP1R " and La Ligue Contre le Cancer N/Ref: RS19/75-127. G.S.B. was supported by a doctoral fellowship from the Fondation de la Recherche Médicale (FRM grant number ECO20170637544) obtained by R.J. D.L. and H.D.P. were supported by funds from the University of Bristol and a project grant from the Biotechnology and Biological Sciences Research Council UK (BBSRC grant number BB/R019223). We thank Lourdes Muñozand Carolina Cera (SimChem, IQAC-CSIC, Barcelona) for support in the synthesis and analysis of compounds.

## ABBREVIATIONS

DAD: diode array detector
DEAC: diethylaminocoumarin
DMAP: 4-dimethylaminopyridine
DMNB: 4,5-dimethoxy-2-nitrobenzyl
FBS: fetal bovine serum
GPCR: G protein-coupled receptor
*o-*NB: *o*-nitrobenzyl
PDA: photodiode array detector
PPG: photocleavable-protecting group
SAR: structure-activity relationship
SCN: suprachiasmatic nucleus
SEM: standard error of the mean
UV-Vis: ultraviolet-visible

---

**Scheme 1.**
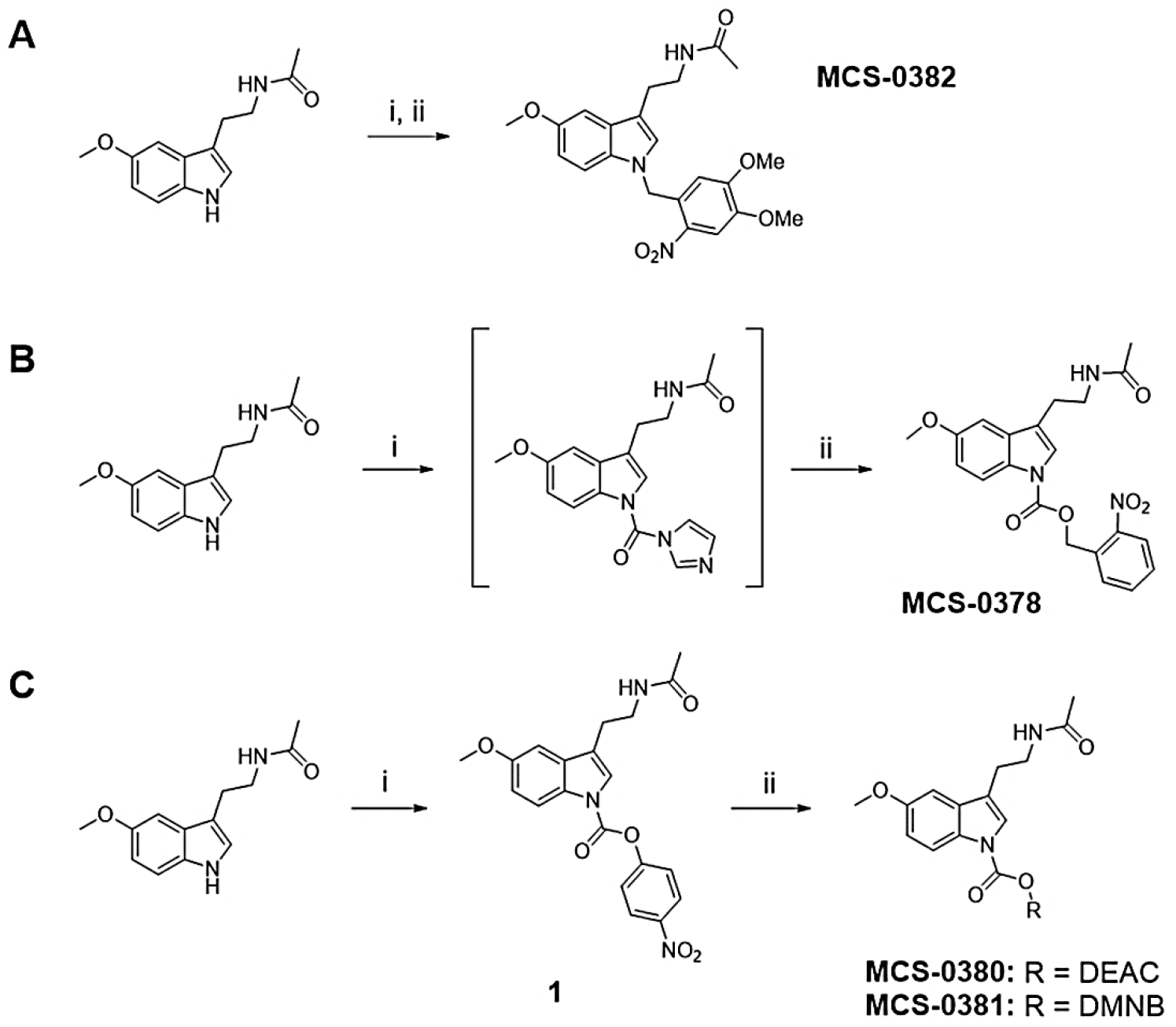
Chemical synthesis of compounds MCS-0378, MCS-0380, MCS-0381 and MCS-0382. Reagents and conditions: (A) i) NaH, DMF, 0 °C, 30 min, ii) DMNB-Br, DMF, -40 °C, 15h, 35%; (B) i) CDI, DMAP, ACN, 40 °C, 30 min, ii) *o*-NB alcohol, ACN, reflux, 2 days, 15%, (C) Reagents and conditions: i) 4-nitrophenyl carbonochloridate, TEA, DMAP, ACN, reflux, 12h, 49%, ii) DEAC-OH or DMNB-OH, DMAP, ACN, reflux, 3 or 6 days, 53% and 26% for **MCS-0380** and **MCS-0381**, respectively Details on the synthesis of DEAC-OH are provided in the Supplementary Information (**Scheme S1**).

