## Additional synthetic procedures, photochemical and pharmacological data for "Design and validation of the first family of photo-activatable ligands for melatonin receptors"

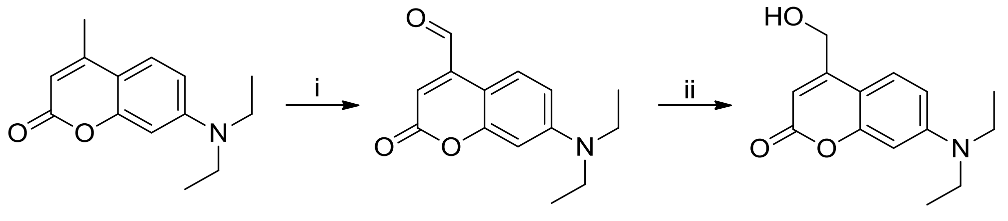


**Scheme S1.** Chemical synthesis of DEAC-OH. Reagents and conditions: **i)** $SeO_2$, Dioxane:Water (15:1), reflux, 2 weeks, 49%; **ii)** $NaBH_4$, EtOH, RT, ON, 11%.

7-(diethylamino)-2-oxo-2H-chromene-4-carbaldehyde **(2)**: to a stirred solution of the commercially available 7-(diethylamino)-4-methyl-2H-chromen-2-one (10.92 g, 46.2 mmol) in a mixture of Dioxane/Water (15:1) (170 mL, 0.25 M), selenium dioxide (12.85 g, 116 mmol) was added. The reaction was heated under reflux for two weeks under magnetic stirring, and the mixture was then filtered through a pad of celite to remove the selenium. The filtered was removed under reduced pressure to yield 7-(diethylamino)-2-oxo-2H-chromene-4-carbaldehyde as a dark red solid (5.22 g, 46%), which was used on the next reaction step without further purification. $^1$H was in good agreement with that described in the literature.[1] $^1$H NMR (400 MHz, Chloroform-*d*): $\delta$ = 10.04 (s, 1H), 8.32 (d, *J* = 9.2 Hz, 1H), 6.66 (dd, *J* = 9.4, 2.5 Hz, 1H), 6.56 (d, *J* = 2.6 Hz, 1H), 6.47 (s, 1H), 3.43 (q, *J* = 7.1 Hz, 4H), 1.22 (t, *J* = 7.1 Hz, 6H).

7-(diethylamino)-4-(hydroxymethyl)-2H-chromen-2-one **(DEAC-OH)**: to an ice-cooled solution of 7-(diethylamino)-2-oxo-2H-chromene-4-carbaldehyde (13 g, 53 mmol, 1 eq) in ethanol (300 mL, 0.18 M), sodium tetrahydroborate (10.44 g, 265 mmol, 5 eq) was added portionwise, and the mixture was stirred overnight at room temperature. Afterwards, the reaction was quenched over HCl 1M (50 mL) and extracted with DCM (4 x 100 mL). Combined organic layers were dried over anhydrous $Na_2SO_4$, filtered and evaporated under reduced pressure. The crude was purified by column chromatography through silica gel using DCM/Acetone (8:2), giving the expected product as a beige solid (1.46 g, 11%). $^1$H was in good agreement with that described in the literature.[1] $^1$H NMR (400 MHz, Chloroform-*d*): $\delta$ = 7.29 (d, *J* = 9.0 Hz, 1H), 6.54 (dd, *J* = 9.0, 2.6 Hz, 1H), 6.45 (d, *J* = 2.7 Hz, 1H), 6.26 (d, *J* = 1.5 Hz, 1H), 4.81 (s, 2H), 3.38 (q, *J* = 7.1 Hz, 4H), 1.18 (t, *J* = 7.1 Hz, 6H). HPLC-PDA-MS (using method B): RT = 3.21 min, $\lambda_{max}$ = 210, 246, 380 nm; purity 97% (254 nm).

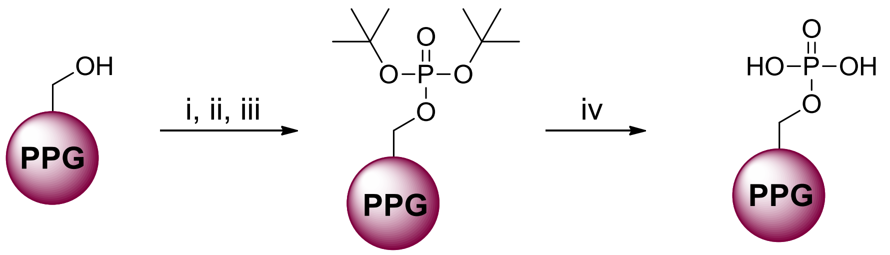

PPG = *o*-NB, **3** DMNB, **4** DEAC, **5**

PPG = *o*-NB, **MCS-1136** DMNB, **MCS-1138** DEAC, **MCS-1140**

**Scheme S2.** Chemical synthesis of the phosphate derivatives MCS-1136, MCS-1138 and MCS-1140. Reagents and conditions: **i)** Tetrazole, DCM, RT, 2 h; **ii)** Di-tert-butyl diethylphosphoramidite, RT, ON; **iii)** TBHP, 0 °C to RT, ON, 32%, 57% and 67% for 3, 4 and 5, respectively; **iv)** HCl, Dioxane, 0 °C to RT, 3 h, 59%, 90% and 92% for MCS-1136, MCS-1138 and MCS-1140, respectively.

**General procedure for the synthesis of the tert-butyl phosphate derivatives 3, 4, 5:** to a solution of the corresponding alcohol (1 eq, 0.5 mmol) in dry DCM (0.1 M), a solution of 1*H*-tetrazole 0.45M (3-4% in wt.) (2 eq) was added, and the reaction was stirred for 2 hours at room temperature. Di-tert-butyl diethylphosphoramidite (1.5 eq) was then added, and the mixture was stirred overnight at room temperature. Afterwards, the reaction was cooled down to 0 ºC and *tert*-butyl hydroperoxide (2.5 eq) was added. The ice/water bath was removed after the addition, and the reaction was stirred until reaction completion at room temperature. The reaction was quenched with an aqueous solution of $Na_2SO_3$ (25 mL) and the aqueous layer was extracted with EtOAc (4 x 25 mL). The organic layers were washed with brine (2 x 50 mL), dried over anhydrous $MgSO_4$, filtered and evaporated under reduced pressure. The crude was purified by column chromatography through silica gel using Hexane/EtOAc as eluents.

Di-tert-butyl 2-nitrobenzyl phosphate **(3)**: this compound was synthesized following the general procedure to obtain tert-butyl phosphate derivatives, starting from (2-nitrophenyl)methanol (89.92 mg, 0.57 mmol), 1H-tetrazole 0.45M (3-4% in wt.) (2.53 mL, 1.14 mmol), di-tert-butyl diethylphosphoramidite (238 µL, 0.85 mmol) and tert-butyl hydroperoxide (259 µL, 1.42 mmol) in 5 mL dry DCM. The reaction was completed three hours after the addition of the tert-butyl hydroperoxide. The crude was purified by column chromatography through silica gel using Hexane/EtOAc (1:1), giving the expected product as a colorless oil (63 mg, 32%). $^1$H-NMR (400 MHz, Chloroform-*d*): δ = 8.14 (d, *J* = 8.1 Hz, 1H), 7.88 (d, *J* = 7.8 Hz, 1H), 7.69 (t, *J* = 7.5 Hz, 1H), 7.48 (t, *J* = 7.7 Hz, 1H), 5.42 (d, *J* = 6.9 Hz, 2H), 1.50 (s, 18H). $^{13}$C NMR (101 MHz, Chloroform-*d*): δ = 146.94, 134.07, 133.75, 128.71, 128.48, 124.95, 83.14, 83.06, 65.30/65.25 (1C), 30.05 (3C), 30.01 (3C). $^{31}$P-NMR (162 MHz, Chloroform-*d*): δ = -10.42. HPLC-PDA-MS (using method B): RT = 3.83 min, $\lambda_{max}$ = 210, 262 nm; purity 94% (254 nm). HRMS calculated for $C_{15}H_{25}NO_6P$: 346.1420 $[M+1]^+$, found: 346.1432.

Di-tert-butyl ((7-(diethylamino)-2-oxo-2*H*-chromen-4-yl)methyl) phosphate **(4)**: this compound was synthesized following the general procedure to obtain tert-butyl phosphate derivatives, starting from DEAC-OH (99.2 mg, 0.39 mmol), 1H-tetrazole 0.45M (3-4% in wt.) (1.73 mL, 0.78 mmol), di-tert-butyl diethylphosphoramidite (166 µL, 0.58 mmol) and tert-butyl hydroperoxide (177 µL, 0.97 mmol) in 5 mL dry DCM. The reaction was completed three hours after the addition of the tert-butyl hydroperoxide. The crude was purified by column chromatography through silica gel using Hexane/EtOAc (2:3), giving the expected product as a yellow oil (96.6 mg, 57%). $^1$H-NMR (400 MHz, Chloroform-*d*): δ = 7.31 (d, *J* = 8.9 Hz, 1H), 6.59 (d, *J* = 8.9 Hz, 1H), 6.53 (s, 1H), 6.25 (s, 1H), 5.10 (d, *J* = 5.7 Hz, 2H), 3.41 (q, *J* = 7.1 Hz, 4H), 1.51 (s, 18H), 1.20 (t, *J* = 7.1 Hz, 6H). $^{13}$C NMR

(101 MHz, Chloroform-*d*): δ = 162.09, 156.28, 150.30, 150.20, 124.53 (3C), 108.96, 106.68, 98.19, 83.45, 83.38, 63.91/63.86 (1C), 45.06, 30.07 (3C), 30.03 (3C), 12.52 (2C). $^{31}$P-NMR (162 MHz, Chloroform-*d*): δ = -9.86. HPLC-PDA-MS (using method B): RT = 4.10 min, $\lambda_{max}$ = 212, 248, 368, 382 nm; purity 95% (254 nm). HRMS calculated for $C_{22}H_{35}NO_6P$: 440.2202 $[M+1]^+$, found: 440.2219.

Di-tert-butyl 4,5-dimethoxy-2-nitrobenzyl phosphate **(5)**: this compound was synthesized following the general procedure to obtain tert-butyl phosphate derivatives, starting from (4,5-dimethoxy-2-nitrophenyl)methanol (131.58 mg, 0.60 mmol), 1H-tetrazole 0.45M (3-4% in wt.) (2.66 mL, 1.20 mmol), di-tert-butyl diethylphosphoramidite (255 µL, 0.90 mmol) and tert-butyl hydroperoxide (272 µL, 1.50 mmol) in 6 mL dry DCM. The reaction was completed three hours after the addition of the tert-butyl hydroperoxide. The crude was purified by column chromatography through silica gel using Hexane/EtOAc (1:1), giving the expected product as a yellow oil (162.5 mg, 67%). $^1$H-NMR (400 MHz, Chloroform-*d*): δ = 7.72 (s, 1H), 7.30 (s, 1H), 5.40 (d, *J* = 6.9 Hz, 2H), 3.98 (s, 3H), 3.94 (s, 3H), 1.50 (s, 18H). $^{13}$C NMR (101 MHz, Chloroform-*d*): δ = 153.97, 147.95, 138.89, 129.19, 109.54, 107.95, 83.10, 83.03, 65.56/65.52 (1C), 56.61, 56.49, 30.01 (3C), 29.97 (3C). $^{31}$P-NMR (162 MHz, Chloroform-*d*): δ = -10.50. HPLC-PDA-MS (using method B): RT = 3.79 min, $\lambda_{max}$ = 238, 305, 340 nm; purity 98% (254 nm). HRMS calculated for $C_{17}H_{29}NO_8P$: 406.1631 $[M+1]^+$, found: 406.1621.

**General procedure to obtain phosphate derivatives MCS-1136, MCS-1138 and MCS-1140:** to a stirred solution of the corresponding tert-butyl phosphate (0.20 mmol, 1 eq) in dioxane (0.1 M), a solution of HCl (50 eq) in dioxane (4M) was added slowly at 0 ºC. The reaction was stirred at room temperature for 3 hours and then evaporated under reduced pressure. The resulting crude was solved in MilliQ water (10 mL) and washed with diethyl ether (3 x 25 mL). The aqueous layer was evaporated under reduced pressure, giving the phosphate derivative without any further purification.

2-nitrobenzyl dihydrogen phosphate **(MCS-1136)**: this compound was obtained following the general procedure 5, starting from di-tert-butyl 2-nitrobenzyl phosphate (72.1 mg, 0.21 mmol) and HCl (2.71 mL, 10.86 mmol) in 2.6 mL dioxane. Evaporation of the reaction mixture after three hours gave the expected product as a white solid (28.7 mg, 59%)**.** $^1$H-NMR (400 MHz, Deuterium Oxide-*d$_2$*): δ = 8.17 (d, *J* = 8.2 Hz, 1H), 7.90 – 7.74 (m, 1H), 7.58 (t, *J* = 7.6 Hz, 1H), 5.33 (d, *J* = 7.6 Hz, 3H). $^{13}$C NMR (101 MHz, Deuterium Oxide-*d$_2$*): δ = 146.44, 134.43, 133.47, 128.60, 128.53, 124.83, 64.26/64.22 (1C). $^{31}$P-NMR (162 MHz, Deuterium Oxide-*d$_2$*): δ = 0.05. HPLC-PDA-MS (using method B): RT = 0.96 min, $\lambda_{max}$ = 210, 268 nm; purity 90% (254 nm). HRMS calculated for $C_7H_9NO_6P$: 234.0168 $[M+1]^+$, found: 234.0166. Melting point: 130.5 - 133.8 ºC

(7-(diethylamino)-2-oxo-2H-chromen-4-yl)methyl dihydrogen phosphate **(MCS-1138)**: this compound was obtained following the general procedure 5, starting from di-tert-butyl ((7-(diethylamino)-2-oxo-2H-chromen-4-yl)methyl) phosphate (96.62 mg, 0.22 mmol) and HCl (2.9 mL, 11.43 mmol) in 2.9 mL dioxane. Evaporation of the reaction mixture after three hours gave the expected product as a yellow oil (64.8 mg, 90%). $^1$H-NMR (400 MHz, Deuterium Oxide-*d$_2$*): δ = 7.97 (d, *J* = 8.7 Hz, 1H), 7.67 (s, 1H), 7.55 (d, *J* = 8.6 Hz, 1H), 6.76 (s, 1H), 5.21 (d, *J* = 7.4 Hz, 2H), 3.72 (q, *J* = 7.2 Hz, 4H), 1.15 (t, *J* = 7.2 Hz, 6H). $^{13}$C NMR (Deuterium Oxide-*d$_2$*): δ = 162.80, 153.51, 152.26, 152.18, 139.31, 126.77, 118.64, 118.40, 113.22, 111.37, 62.62, 62.58, 53.72, 9.55. $^{31}$P-NMR (162 MHz, Deuterium Oxide-*d$_2$*): δ = 0.03. HPLC-PDA-MS (using method B): RT = 2.69 min, $\lambda_{max}$ = 210, 250, 386 nm; purity > 98% (254 nm). HRMS calculated for $C_{14}H_{19}NO_6P$: 382.0950 $[M+1]^+$, found: 382.0959 .

4,5-dimethoxy-2-nitrobenzyl dihydrogen phosphate **(MCS-1140)**: this compound was obtained following the general procedure 5, starting from di-tert-butyl 4,5-dimethoxy-2-nitrobenzyl phosphate (143.10 mg, 0.35 mmol) and HCl 4M (4.6 mL, 18.36 mmol) in 4.5 mL dioxane. Evaporation of the reaction mixture after three hours gave the expected product as a yellow oil (94.8 mg, 92%). $^1$H-NMR (400 MHz, Deuterium Oxide-*d$_2$*): δ = 7.82 (s, 1H), 7.38 (s, 1H), 5.29 (d, *J* = 7.4 Hz, 2H), 4.01 (s, 3H), 3.94 (s, 3H). $^{13}$C NMR (101 MHz, Deuterium Oxide-*d$_2$*): δ = 153.46, 147.05, 139.95, 129.94, 110.00, 108.39, 64.26, 56.22, 56.05. $^{31}$P-NMR (162 MHz, Deuterium Oxide-*d$_2$*): δ = 0.11. HPLC-PDA-MS (using method B): RT = 2.28 min, $\lambda_{max}$ = 216, 242, 350 nm; purity > 98% (254 nm). HRMS calculated for $C_9H_{13}NO_8P$: 294.0379 $[M+1]^+$, found: 294.0385. Melting point: 138.2-138.3 ºC

#### Additional photochemical data

**Table S1.** Affinity of melatonin, AFMK, MCS-1136, MCS-1138 and MCS-1140 for $MT_1$ and $MT_2$

| Ligand | $pK_i$ ± S.E.M. (M) | |
|---|---|---|
| | $MT_1$ | $MT_2$ |
| Melatonin | 9.56 ± 0.29 | 9.76 ± 0.45 |
| Melatonin + Light | 9.43 ± 0.19 | 9.57 ± 0.16 |
| AFMK | 5.37 ± 0.48 | 5.67 ± 0.09 |
| AFMK + Light | 4.94 ± 0.97 | 6.05 ± 0.07 |
| MCS-1136 | n.d. | n.d. |
| MCS-1136 + Light | n.d. | n.d. |
| MCS-1138 | n.d. | n.d. |
| MCS-1138 + Light | n.d. | 6.15 ± 0.48 |
| MCS-1140 | n.d. | n.d. |
| MCS-1140 + Light | n.d. | n.d. |

The affinity was measured in 2-[$^{125}$I]-MLT competition experiments and is expressed as mean $pK_i$ ± S.E.M. (M). Data correspond to the mean of at least three independent experiments, each of them performed using at least eight different ligand concentrations. Values obtained with light-activated compounds correspond to apparent $pK_i$.

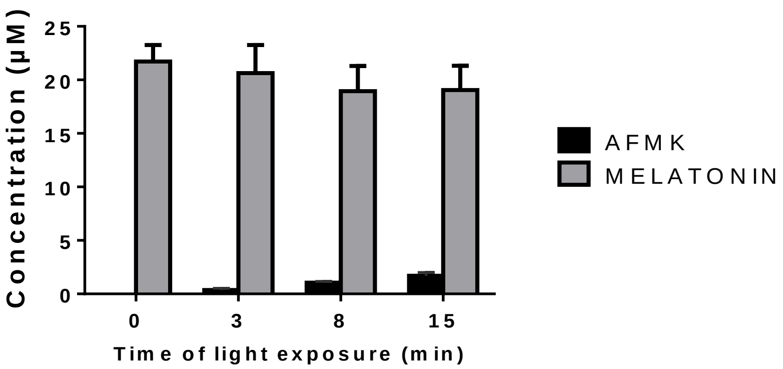


**Figure S1.** Quantification of melatonin and AFMK concentrations upon irradiation of a 20 μM solution of melatonin in aqueous buffer, after applying light for different periods of time (t = 0, 3, 8 and 15 min). Light is applied to the ligands for 10 minutes prior to incubation with the cells, using the laser as a light source (405 nm, 100 mW). Data are expressed as mean ± SEM of three independent experiments

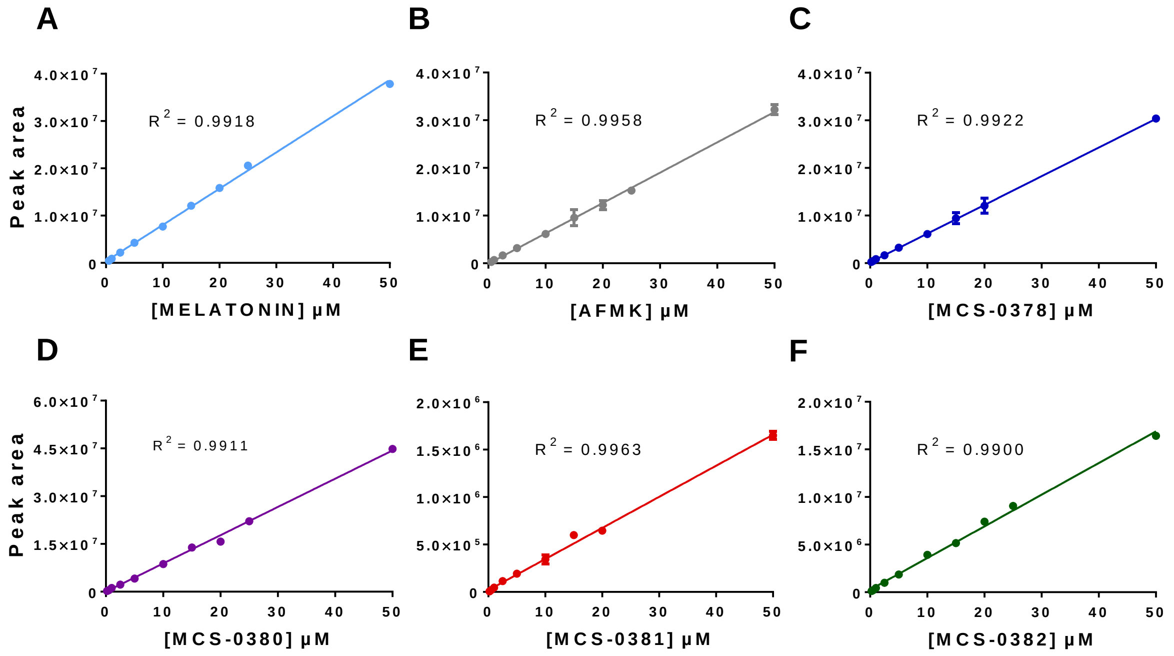


**Figure S2.** Calibration curves of melatonin **(A)**, AFMK **(B)**, MCS-0378 **(C)**, MCS-0380 **(D)**, MCS-0381 **(E)** and MCS-0382 **(F)**. Concentrations (µM) are plotted against peak area with least-squares linear regression. Data are expressed as mean of the peak area ± S.E.M. of at least three independent experiments.

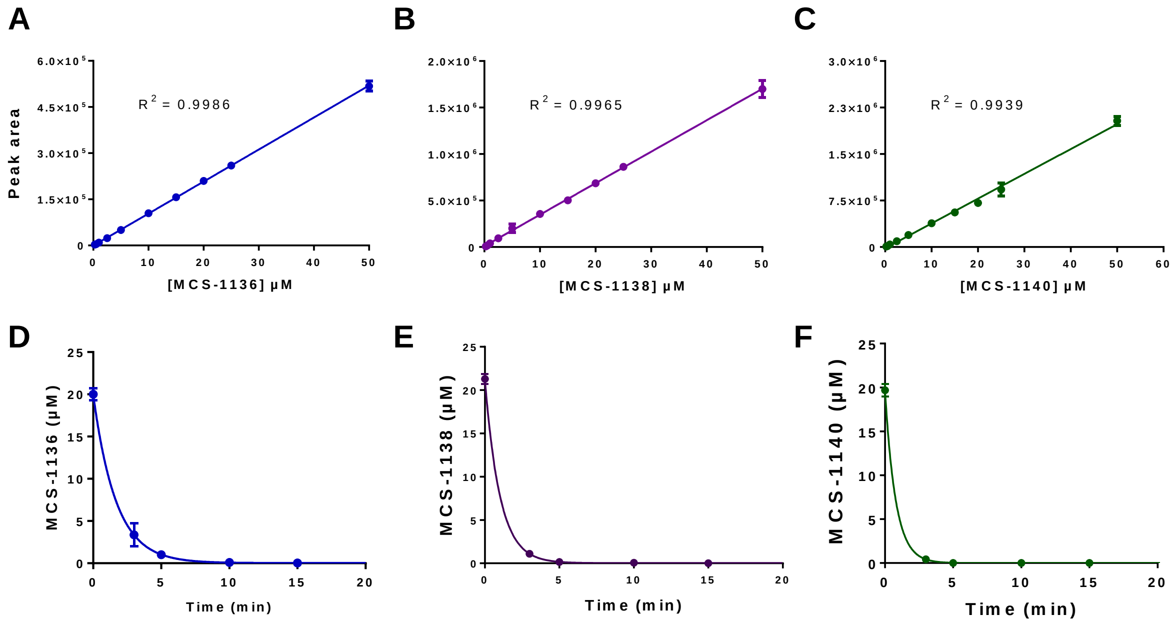


**Figure S3.** Photolytic assays of the phosphate-derivatives MCS-1136, MCS-1138 and MCS-1140. Calibration curves of compounds MCS-1136 **(A)**, MCS-1138 **(B)** and MCS-1140 **(C)**. Concentrations (µM) are plotted against peak area with least-squares linear regression. Data are expressed as mean of the peak area ± S.E.M. of at least three independent experiments. Uncaging rate of MCS-1136 **(D)**, MCS-1138 **(E)** and MCS-1140 **(F)**, expressed as mean of the phosphate concentration ± S.E.M (μM) of at least three independent experiments. Light is applied to the ligands for 10 minutes prior to incubation with the cells, using the laser as a light source (405 nm, 100 mW).

### Additional pharmacological data

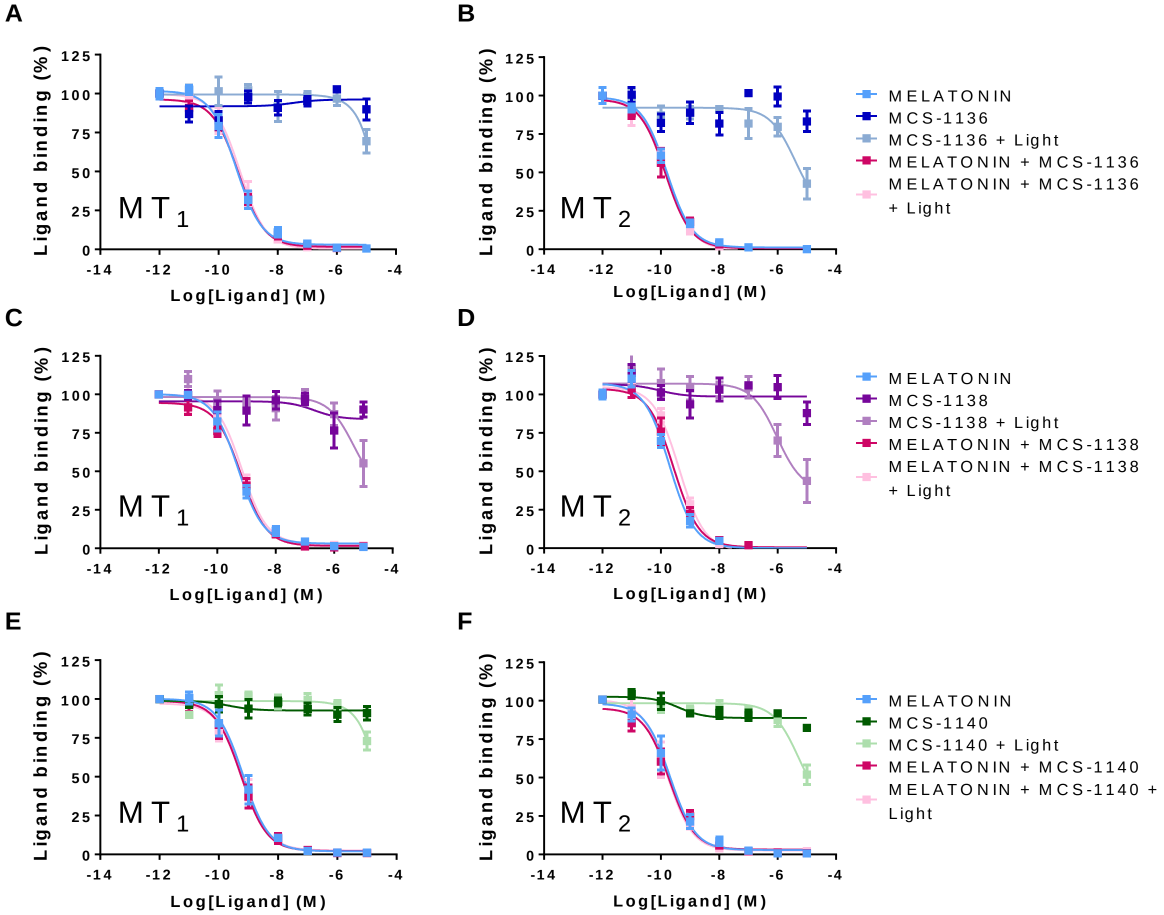


**Figure S4.** Competition of 2-[$^{125}$I]-MLT binding by melatonin and compounds MCS-1136 **(A, B)**, MCS-1138 **(C, D)** and MCS-1140 **(E, F)** before and after light exposure, in crude membranes from HEK293 cells expressing human $MT_1$ or $MT_2$, respectively. Light is applied to the ligands for 10 minutes prior to incubation with the cells, using the laser as a light source (405 nm, 100 mW). Data are expressed as mean ± SEM of three independent experiments. Data are represented as percentage of maximal binding in the absence of compounds and normalized to melatonin maximum effect (= 100%).

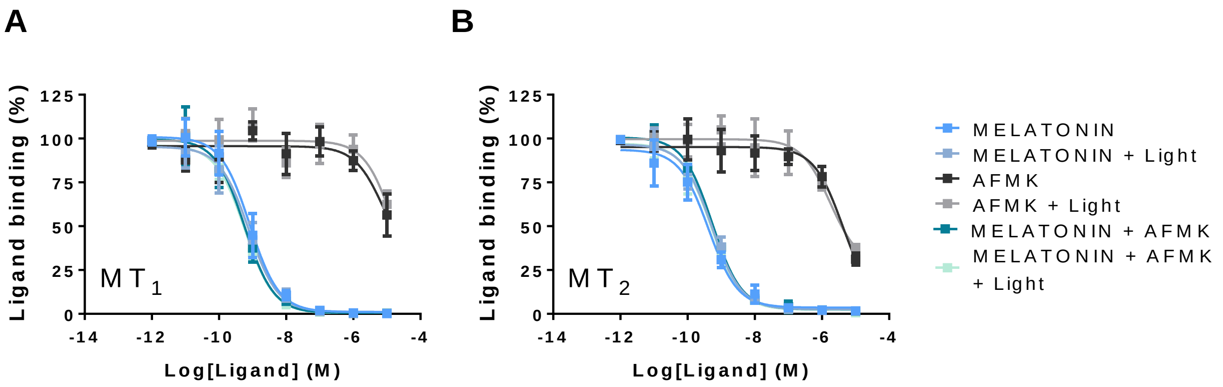


**Figure S5.** Competition of 2-[$^{125}$I]-MLT binding by melatonin and AFMK before and after light exposure, in crude membranes from HEK293 cells expressing human $MT_1$ (A) or $MT_2$ (B). Light is applied to the ligands for 10 minutes prior to incubation with the cells, using the laser as a light source (405 nm, 100 mW). Data are expressed as mean ± SEM of three independent experiments. Data are represented as percentage of maximal binding in the absence of compounds and normalized to melatonin maximum effect.
